# Mimicking Kidney Flow Shear Efficiently Induces Aggregation of LECT2, a Protein Involved in Renal Amyloidosis

**DOI:** 10.1101/2023.07.13.548788

**Authors:** Jeung-Hoi Ha, Yikang Xu, Harsimranjit Sekhon, Stephan Wilkens, Dacheng Ren, Stewart N. Loh

## Abstract

Aggregation of leukocyte cell-derived chemotaxin 2 (LECT2) causes ALECT2, a systemic amyloidosis that affects the kidney and liver. Homozygosity of the I40V LECT2 mutation is believed to be necessary but not sufficient for the disease. Previous studies established that LECT2 fibrillogenesis is greatly accelerated by loss of its single bound zinc ion and stirring or shaking. These forms of agitation are often used to facilitate protein aggregation, but they create heterogeneous shear conditions, including air-liquid interfaces that denature proteins, that are not present in the body. Here, we determined the extent to which a more physiological form of mechanical stress—shear generated by fluid flow through a network of artery and capillary-sized channels—drives LECT2 fibrillogenesis. To mimic blood flow through the human kidney, where LECT2 and other proteins form amyloid deposits, we developed a microfluidic device consisting of progressively branched channels narrowing from 5 mm to 20 μm in width. Flow shear was particularly pronounced at the branch points and in the smallest capillaries, and this induced LECT2 aggregation much more efficiently than conventional shaking methods. EM images suggested the resulting fibril structures were different in the two conditions. Importantly, results from the microfluidic device showed the first evidence that the I40V mutation accelerated fibril formation and increased both size and density of the aggregates. These findings suggest that kidney-like flow shear, in combination with zinc loss, acts in combination with the I40V mutation to trigger LECT2 amyloidogenesis. These microfluidic devices may be of general use for uncovering the mechanisms by which blood flow induces misfolding and amyloidosis of circulating proteins.

## Introduction

Since its initial report in 2008^1^, systemic amyloidosis of LECT2 (ALECT2) has risen to become the third most common type of renal amyloidosis in the U.S.^2^. It results from misfolding of leukocyte cell-derived chemotaxin-2 (LECT2), a 133-amino acid protein synthesized in hepatocytes for secretion into the blood. As in all systemic amyloid diseases, the protein misfolds and aggregates in the extracellular space, and in the case of ALECT2 the amyloid fibrils accumulate in the kidneys and liver, leading to their eventual failure. All ALECT2 patients examined to date are homozygous for the I40V mutation^3, 4^ with the exception of one heterozygous individual^5^. The I40V polymorphism, however, approaches 50% allele frequency in certain populations, which has led to consensus in the field that one or more additional conditions are required for ALECT2^3, 6^.

We recently established that loss of LECT2’s single bound zinc ion appears to be obligatory for fibrillogenesis^7^. Zinc helps to maintain the structure and stability of LECT2: removing the metal caused widespread changes in NMR spectra and decreased the midpoints of thermal and guanidinium chloride denaturation. Zinc loss may occur naturally in the body. Measurements of the zinc dissociation constant indicated that a significant percentage of LECT2 exists in the zinc-free form (apoLECT2) at the normal blood pH of 7.4, with this value increasing considerably at pH 6.5. In addition to zinc loss, mechanical agitation by magnetic stirring or shaking with silica beads was found to induce fibril formation. The effects of the I40V mutation on the above properties, however, were subtle. The mutation slightly destabilized apoLECT2 but did not alter holo protein stability, zinc binding affinity, or zinc dissociation rate. The I40V mutation did not consistently accelerate aggregation of apoLECT2 as determined by thioflavin T (ThT) fluorescence in the presence of stirring or shaking, but instead introduced variation in the lag times and amplitudes of aggregation. Thus, the role of the I40V mutation in the disease process remained undefined.

The above results suggested a model in which two perturbants, zinc loss and agitation-induced shear, destabilize and distort (respectively) the LECT2 structure such that a buried amyloidogenic sequence (e.g., residues 52 – 58 or 79 – 88)^8^ becomes transiently exposed. Hydrodynamic shear has been shown to induce amyloid formation of Alzheimer Aβ peptides^9–12^, β-lactoglobulin^13–15^, insulin^15–19^, α-synuclein^20–22^, apolipoprotein C-II^23^, superoxide dismutase^24^, antibodies^25^, and other proteins. Magnetic stirring, shaking with beads, vortexing, and sonication are commonly used for this purpose. These methods produce heterogeneous shear conditions and incur extensive aeration and bubbles^21, 24, 26^, which efficiently denature proteins and induce fibril formation^22, 23^. However, such air-liquid interfaces are not present in the circulatory system.

The main form of mechanical stress experienced by circulating proteins is fluid shear in the absence of an air-liquid interface^29, 30^. Shear is generated by the velocity gradient of fluids traveling through blood vessels and is determined by flow rate, vessel geometry, and viscosity of the blood. This stress can be reproduced by devices in which solutions are passed through small, closed channels, resulting in laminar flow profiles like those typically present in blood vessels^25, 31^. These microfluidic devices have been employed to study the properties and formation of amyloid fibrils. Fibrillization of Alzheimer Aβ42^32, 33^ and insulin^34, 35^ peptides were found to be enhanced under flow compared to bulk conditions.

Here, we developed the first microfluidic device designed to recapitulate the structures and shear forces present in the human kidney, to study aggregation of LECT2 and other proteins involved in systemic amyloidosis. The kidney is highly vascularized with increasingly branched and narrowing vessels which, together with the high blood flow through the organ (∼8 L/h), create flow shear conditions that are extensive and unique. Our findings indicated that microfluidic flow induced amyloid formation much more efficiently than conventional stirring or shaking. Moreover, the data showed for the first time that the I40V mutation caused LECT2 to aggregate more rapidly and into larger, denser bodies. These results suggest that kidney-like flow shear and zinc loss act in concert with the I40V mutation to drive ALECT2.

## Results

### LECT2 stability and zinc binding

Recombinant human LECT2 was prepared as described in our earlier work^7^ with one difference. We removed extraneous amino acids (AAA from the N-terminus and GSGGLEVLFQ from the C-terminus) that were a legacy of the protease sites used to cleave away the fusion protein (ribose binding protein) that was necessary for *E. coli* expression and purification. The protein containing those extra sequences is referred to as LECT2-tag. The protein used in the present study (simply designated as LECT2) is composed of the human sequence with an additional GPG at the N-terminus, without which it was not feasible to cleave away the ribose binding protein.

The zinc dissociation constant (K_d_) of LECT2 was lower than that of LECT2-tag by 4 – 5-fold, with higher affinity mostly attributable to a slower k_off_ (Table 1 and Supporting Figures S1A and S1B). Removing the tag slightly increased thermodynamic stability as determined by guanidine hydrochloride (GdnHCl) denaturation experiments (Table 1 and Supporting Figure S1C). Despite LECT2 being more robustly native than LECT2-tag, the effects of mutation that we reported in our previous study of the latter remained true for the former; namely, (i) the I40V mutation did not alter zinc binding affinity or dissociation rate, and (ii) the I40V mutation slightly destabilized the proteins in their apo forms but not in their holo forms.

**Table 1.** Zinc binding and stability of LECT2. Conditions were 2.5 µM protein, 20 mM Tris(pH 7.5), 0.15 M NaCl, 37 °C. Holo LECT2 samples contained 2.5 µM ZnCl_2_ and 7.5 µM iminodiacetic acid. Errors are s.d. (n = 3 – 4). NA, not applicable. ^a^Ha et al. (2021)^7^. ^b^22 °C.

| **Table 1. Zinc binding and stability of LECT2.** Conditions were 2.5 μM protein, 20 mM Tris (pH 7.5), 0.15 M NaCl, 37 °C. Holo LECT2 samples contained 2.5 μM ZnCl<sub>2</sub> and 7.5 μM iminodiacetic acid. Errors are s.d. (n = 3 – 4). NA, not applicable. <sup>a</sup>Ha et al. (2021)<sup>7</sup>. <sup>b</sup>22 °C. | | | |
| --- | --- | --- | --- |
| **Protein** | **K<sub>d</sub> (nM)** | **k<sub>off</sub> (s<sup>-1</sup>)** | **ΔG<sub>fold</sub> (kcal/mol)** |
| WT LECT2 | 0.14 ± 0.03 | 0.0032 ± 0.0002 | -6.31 ± 0.49 |
| I40V LECT2 | 0.21 ± 0.03 | 0.0025 ± 0.0002 | -6.69 ± 0.12 |
| WT apoLECT2 | NA | NA | -3.35 ± 0.52 |
| I40V apoLECT2 | NA | NA | -3.04 ± 0.08 |
| WT LECT2-tag<sup>a</sup> | 0.79 ± 0.45<sup>b</sup> | 0.0098 ± 0.0021 | -4.16 ± 0.10 |
| I40V LECT2-tag<sup>a</sup> | 0.76 ± 0.27<sup>b</sup> | 0.0093 ± 0.0010 | -3.90 ± 0.006 |
| WT apoLECT2-tag<sup>a</sup> | NA | NA | -2.65 ± 0.05 |
| I40V apoLECT2-tag<sup>a</sup> | NA | NA | -2.18 ± 0.02 |

### Microfluidic shear efficiently induced aggregation of apoLECT2

We modeled the device after the dimensions and branched structure of the renal vasculature (Figure 1A). Blood enters the human kidney through the renal artery (∼5 mm dia.), where it divides into progressively narrower arteries (segmental, interlobular, arcuate, and cortical radiate) and finally into afferent/efferent arterioles (15 – 20 μm dia.) that enter and exit the glomerulus. Microfluidic devices were fabricated using soft lithography in which polydimethylsulfoxane (PDMS) stamps were made from silicon wafer molds. The pattern consisted of a single 5.12 mm wide entry channel that bifurcated eight times into widths of 2.56 mm (x2), 1.28 mm (x4), 640 μm (x8), 320 μm (x16), 160 μm (x32), 80 μm (x64), 40 μm (x128), and 20 μm (x256), whereupon the bifurcation pattern reversed into a single 5.12 mm exit channel. All channels were rectangular and 20 μm high. Microbore tubes were inserted into the inlet and outlet channels and sealed with PDMS. Protein solutions were circulated from reservoirs (sealed and maintained at constant temperature) by means of a peristaltic pump to mimic the pulsatile delivery by the heart as well as enable low-shear pumping action. The PDMS block was mounted on a microscope slide and aggregation was monitored by light microscopy (Figure 1B).

**Figure 1.**
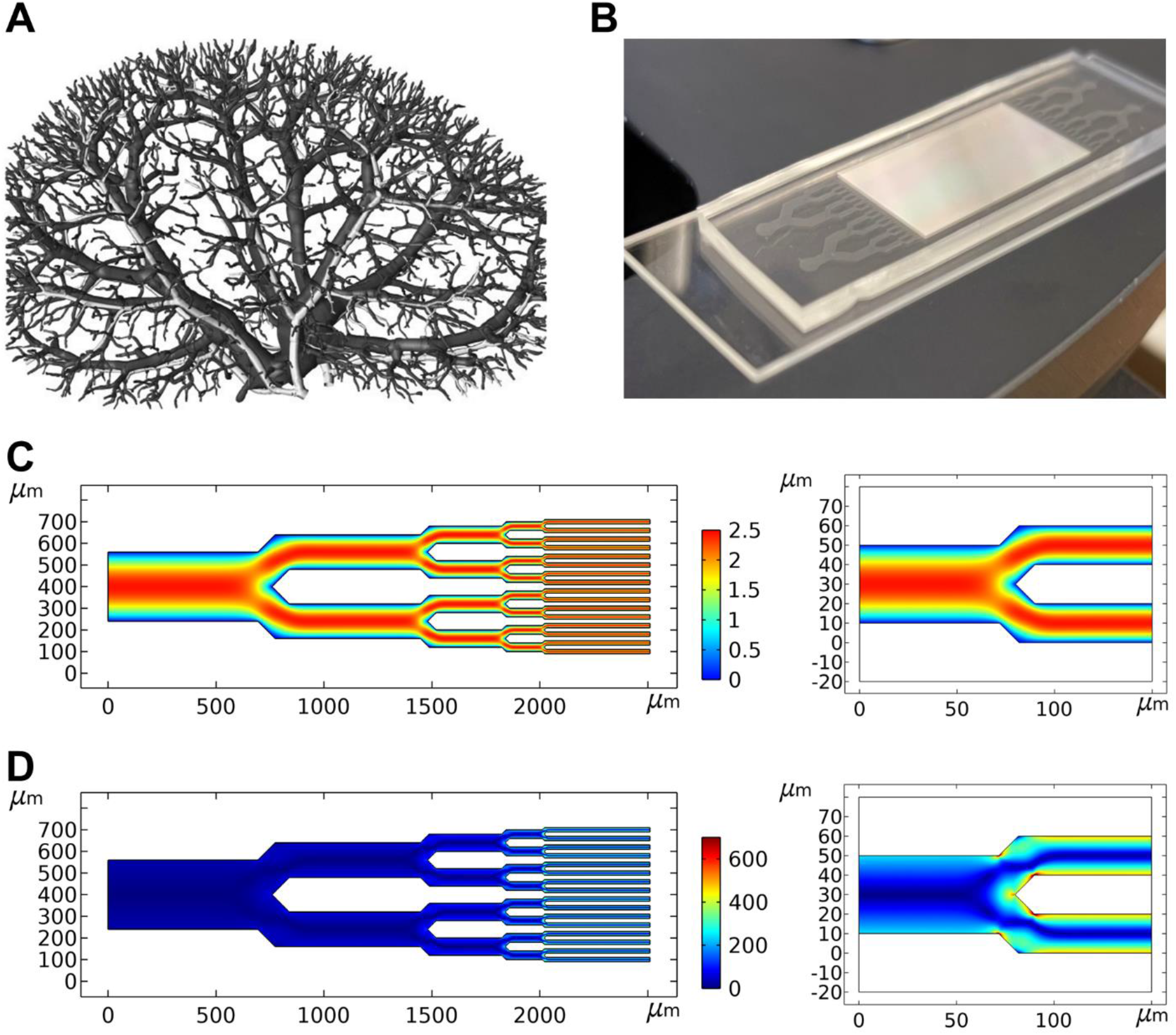
Microfluidic chip design. **(A)** The microfluidic device was designed to mimic the branched vasculature of the human kidney, shown reconstructed from 20 µm micro-CT data^36^. Image reproduced with permission from the publisher. **(B)** I40V and WT LECT2 samples were analyzed in parallel on the same chip. **(C)** Simulations of fluid flow through a 320 µm – 20 µm pathway of channels mapped gradients in flow velocity, revealing areas of potential shear and where flow velocity dropped to nearly zero (cyan/blue; units in mm/s). Velocity gradients were particularly pronounced at the 40/20 µm junctions (right). **(D)** Calculations of shear rate identified regions of shear stress (cyan/yellow/orange; units in 1/s), which were present at all junction points but most evident at the smallest junctions and in the 20 µm channels (right).

The pumping rate was set to 10 μl/min to make the flow rate at each of the 256 narrowest channels in our device (0.04 μl/min) approximately equal to that at the afferent/efferent arterioles (0.05 – 0.1 μl/min). We designed the device so that the average linear flow velocity (1.63 mm/s) and pressure gradient (∼9 mm Hg) stayed constant across all hierarchical dimensions. This allowed us to model shear stress accurately using computational fluid dynamic simulations and determine the effect of branching in the absence of other variables. These simulations confirmed that flow was laminar in all channels (Figure 1C) with Reynolds number well below 1. As such, the principal shear stress arose from the velocity gradient perpendicular to the direction of travel, which became increasingly pronounced as channel width decreased. The junctions introduced additional velocity gradients that caused the fluid traveling in the middle of the channel to slow down as it approached the split point then accelerate as it passed, as shown by the red-orange-red path at the 40/20 µm junction (Figure 1C, right). These gradients also established a ‘dead zone’ where velocity dropped to nearly zero at the tip of the junction. Calculations of shear rate from these data mapped the locations of shear fields, which were most pronounced at the branch points, especially the smaller ones, and in the 20 µm channels themselves (Figure 1D, right). Extensional flow is formed by velocity gradients in the same direction as travel and strain caused by extensional stress can also denature proteins^25, 37^. Calculations indicated that strain was minimal in our device (Supporting Figure S2), confirming that the main stress came from shear. In summary, the computational fluid dynamic simulations suggested that the junctions may facilitate aggregation by providing both elevated shear stress and areas of low velocity flow, where the transiently misfolded proteins can potentially associate before they are able to refold.

At 37 °C and an apoLECT2 concentration of 10 µM we observed aggregation in less than 24 h. Aggregation was highly dependent on temperature, and to lengthen the onset of aggregation to a more observable 24 – 48 h we decreased the temperature of microfluidic experiments to 35 °C. As a negative control, we repeated the experiments using holoLECT2. No aggregation was observed over 3 d, confirming that zinc loss was required for aggregation under microfluidic flow conditions.

ApoLECT2 aggregates tended to form initially at channel bifurcations, particularly at the narrowest (40/20 µm) junctions as predicted by the simulations (**Error! Reference source not found.** and Supporting Figure S3). Deposits then clogged many of the 20 µm channels and accumulated in the upstream channels (Supporting Figure S4). This behavior appeared to resemble the end-stage renal pathology observed in patients with severe amyloidosis, and is consistent with the finding that renal amyloidosis is a disease of small vessels and capillaries^38^. Light microscopy images suggested nucleation-growth kinetics typical of amyloid. The first, small aggregates became visible after 24 – 48 h with large-scale channel blockage occurring shortly thereafter. Rapid growth was likely exacerbated by the shear stress being inversely proportional to the cross-sectional area of the channel, creating a positive feedback loop between the size of the clog and its growth rate.

For quantifying aggregation, we imaged the three narrowest junctions on the inlet side of the chip (160/80 µm, 80/40 µm, and 40/20 µm). One microscope image contained either two 160/80 µm junctions, three 80/40 µm junctions, or six 40/20 µm junctions. Five images of each junction type were recorded per biological repeat (BR; defined as samples prepared independently on different days, frequently from different protein stocks and different fabrication runs of microfluidic chips), accounting for 24 % of the number of junctions in the upstream half of the chip. Representative results from one BR are shown in Figure 3; data from 7 additional BRs are included as Supporting Figures S5 – S11.

To determine the effects of the I40V mutation using the microfluidic device, we first calculated the fraction of channel area that was occupied by particles large enough to block light. The dark pixels in the channels were counted and this value was divided by the total number of channel pixels. These fractional areas confirmed the visual impression from Figure 2 that I40V apoLECT2 formed larger deposits than WT at most time points and junction sizes (Figure 3, left plots). The differences ranged from 2.5-fold to 6.8-fold for the cases where p-values indicated the highest confidence (p<10^-3^).

**Figure 2.**
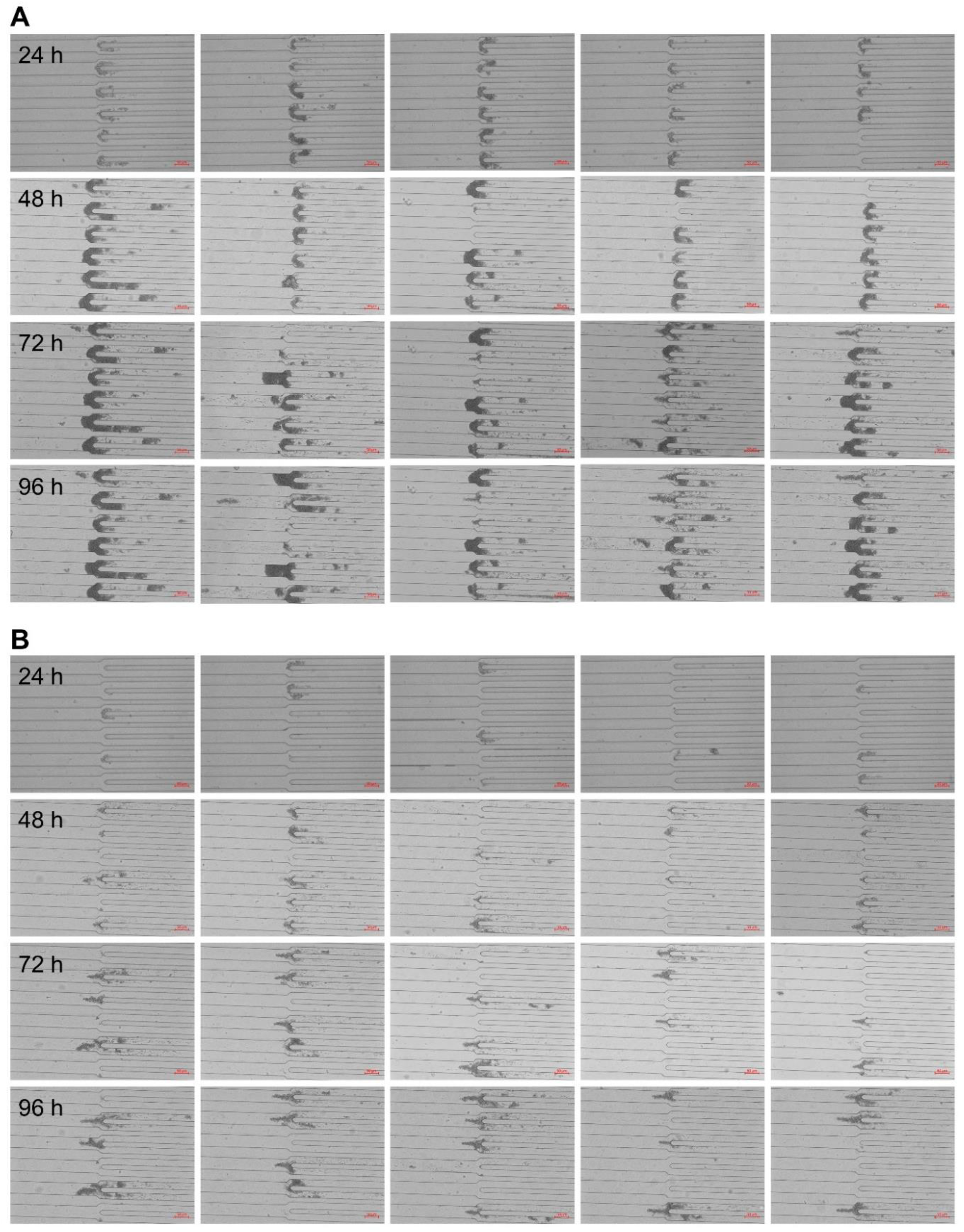
Microfluidic device efficiently induced aggregation of I40V apoLECT2. Images of the narrowest junctions (40/20 µm) are shown for I40V **(A)** and WT **(B)** apoLECT2. Scale bars, 50 μm. See Supporting Figures S3 and S4 for additional images.

**Figure 3.**
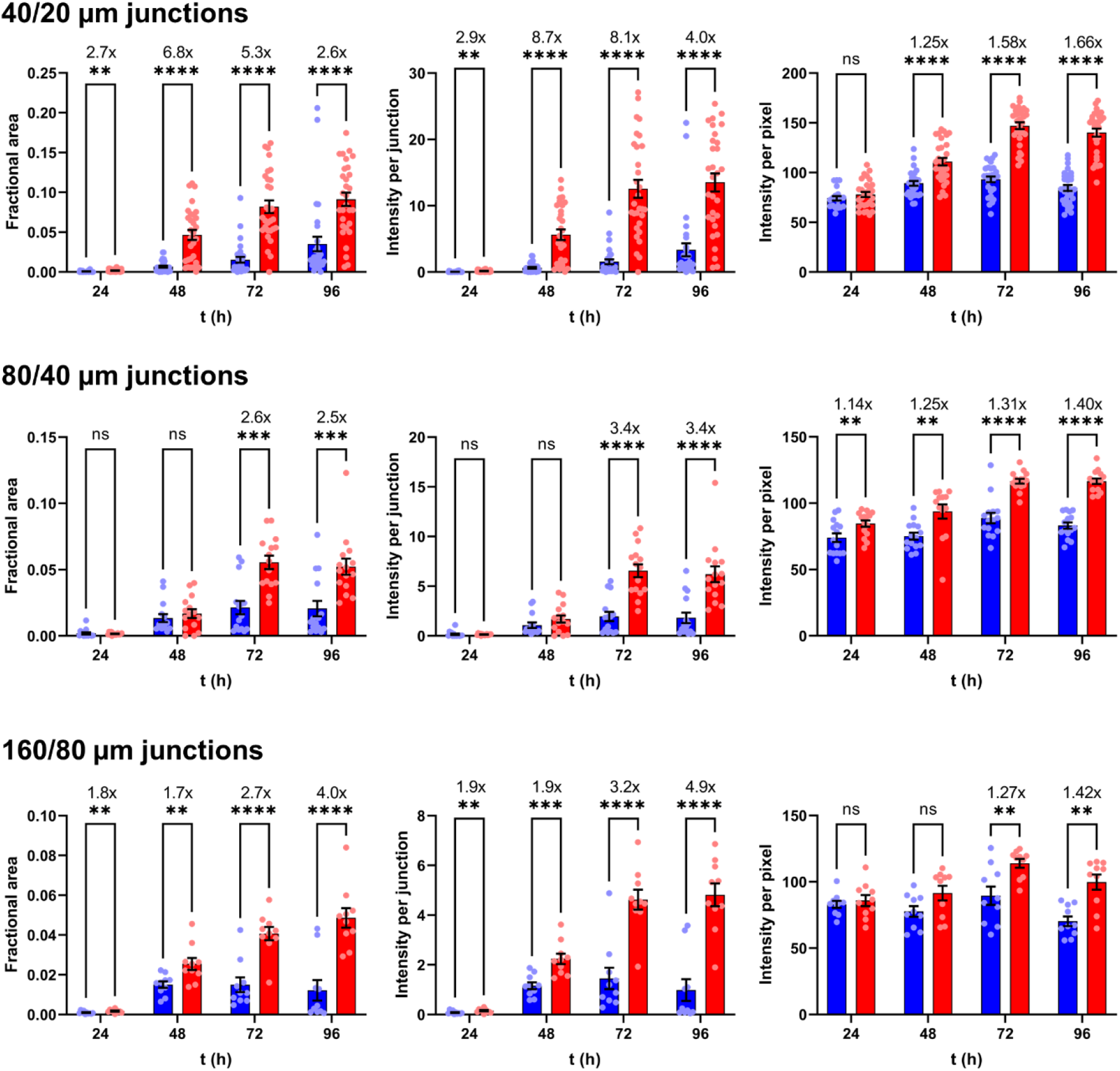
The I40V mutation increased size and density of aggregates induced by flow shear. Red and blue data indicate I40V and WT apoLECT2, respectively. Sizes of deposits were characterized by their fractional areas (left plots), and densities of deposits by their integrated intensities (center plots) and per-pixel intensities (right plots). Fold changes between I40V and WT, when significant, are indicated by the numbers above the bars. The number of data points (circles) for the 40/20 µm, 80/40 µm, and 160/80 µm junctions were 30, 15, and 10, respectively. ****, p<10^-4^; ***, p<10^-3^; **, p<10^-2^; ns, not significant. See Supporting Figures S5 – S11 for additional BR data.

We then quantified the density of the aggregates by measuring the intensities of the same dark pixels that were identified in the fractional area analysis above (a representative heat map image is shown in Figure 4A), summing these values, subtracting the average pixel intensity of empty channels, and dividing by the total number of channel pixels. As expected, the integrated intensities (Figure 3, middle plots) showed the same trend as the fractional areas (Figure 3, left plots), but the fold change between I40V and WT increased at all time points and channel sizes, suggesting that the I40V aggregates were denser. This conclusion was supported by determining the average intensity per pixel of aggregate (Figure 3, right plots). The per-pixel intensity data were much more tightly grouped compared to the fractional area and integrated intensity data. The reason is that the channels displayed a wide distribution of aggregate sizes (including zero), and therefore a wide range of integrated intensities, but dividing the latter values by the former (excluding channels with zero aggregates) removed this variability. The increased statistical confidence of the per-pixel intensity analysis allowed us to conclude that the I40V aggregates were denser than the WT aggregates at all time points, with the difference becoming more pronounced at longer times. Together, these data indicate that the I40V mutation caused apoLECT2 to aggregate faster and into larger, denser bodies under flow shear conditions.

**Figure 4.**
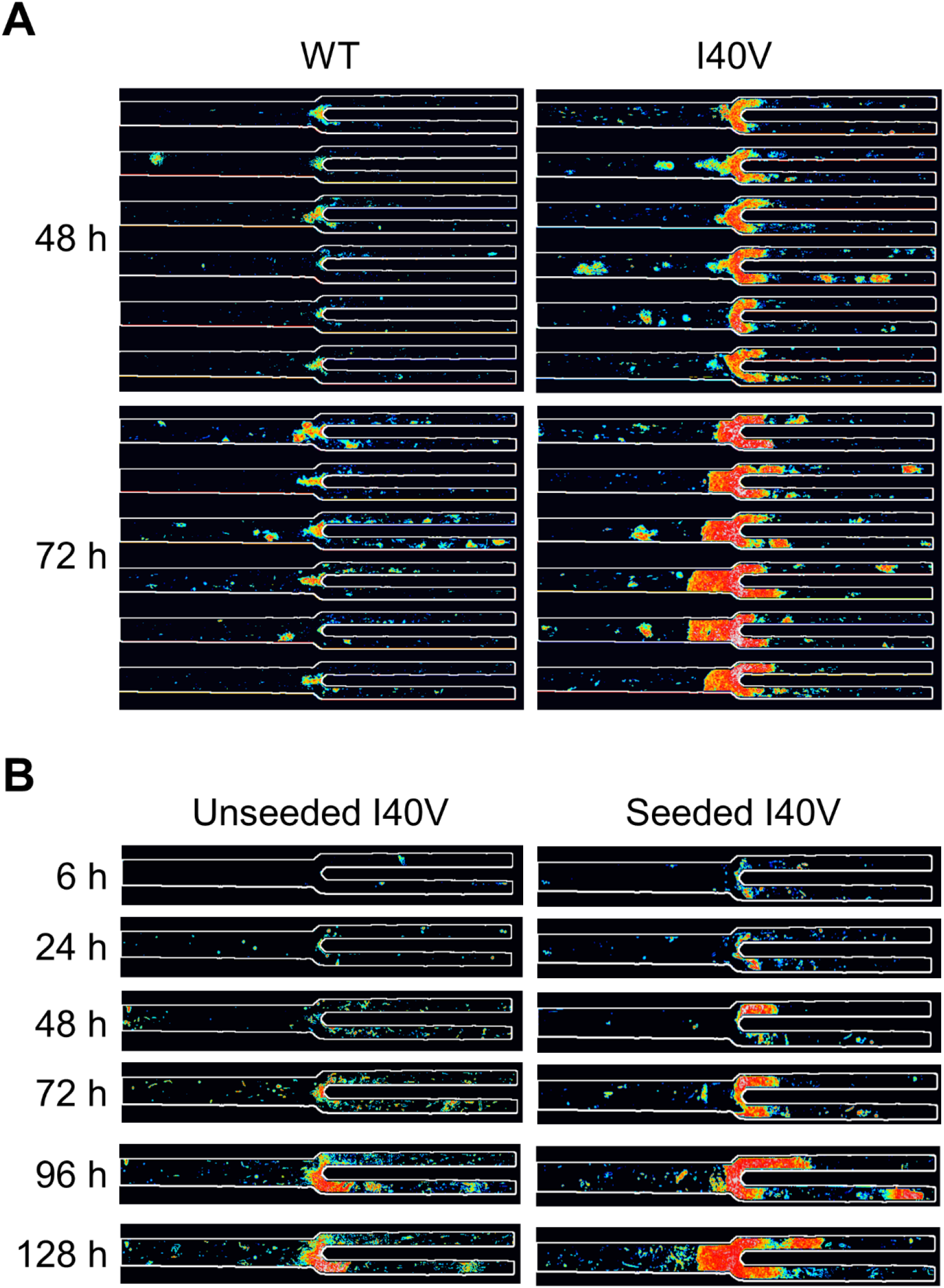
Characterization of size and density of aggregates formed by flow shear. **(A)** Deposits of I40V apoLECT2 were larger and denser than those of WT apoLECT2 as determined by measuring pixel intensities and representing them as a pseudocolored heat map (blue is least intense and red is most intense). **(B)** Seeding decreased the lag time of I40V apoLECT2 aggregation. Each image shown is the sum of five individual images of 40/20 µm junctions.

### Shaking-induced aggregation

To compare the microfluidic results to those obtained by conventional methods, we monitored protein aggregation by shaking the samples with 1 mm silica beads in the presence of thioflavin T (ThT), a dye that becomes more fluorescent upon binding to amyloid fibrils. LECT2 aggregation was chaotic under these conditions, as we reported previously^7^, with lag phases and fluorescence amplitudes varying considerably among BRs and even among technical replicates (TR; defined as samples from the same BR divided into multiple wells in the microtiter plate). The source of variability was unclear but was likely due to the inherently heterogeneous nature of aeration/agitation exacerbated by noticeable differences in bead shape. For this reason, we show raw data for all replicates without averaging or normalization.

Like in the flow shear experiments, I40V apoLECT2 aggregated more aggressively compared to WT under shaking conditions. Thirty-five percent of the 31 replicates of I40V apoLECT2 exhibited increased ThT fluorescence over the 160 – 400 h duration of the experiments, whereas all 32 WT apoLECT2 samples remained at baseline (Supporting Figure S12A and Table 2). Typical lag times of I40V apoLECT2 were 80 – 100 h. These results differ from those of our earlier apoLECT2-tag study, which found that both I40V and WT apoLECT2- tag aggregated rapidly and with indistinguishable lag phases (5 – 10 h)^7^. This discrepancy may be due to the increased stability of the tag-free proteins (Table 1). The data suggest that slowing down the overall rate of aggregation may have allowed the molecular differences between I40V and WT apoLECT2 to manifest in the shaking experiments. That the trends in shaking results corroborate those seen in microfluidic experiments demonstrates that the I40V mutation enhances aggregation propensity of apoLECT2 in different mechanical stress conditions.

**Table 2.** Number of apoLECT2 samples that exhibited increased ThT fluorescence in shaking experiments. Raw data and the number of BR and TR are shown in Supporting Figure S3. Conditions are 10 μM apoLECT2, 5 μg/ml HS, 5 μg/ml heparin, 34 °C.

|  |
| --- |
| **Table 2. Number of apoLECT2 samples that exhibited increased ThT fluorescence in shaking experiments.** Raw data and the number of BR and TR are shown in Supporting Figure S3. Conditions are 10 μM apoLECT2, 5 μg/ml HS, 5 μg/ml heparin, 34 °C. |

| ApoLECT2 | No additive | HS | Heparin | HoloLECT2 |
| --- | --- | --- | --- | --- |
| I40V | 11/31 | 34/39 | 25/27 | 0/18 |
| WT | 0/32 | 7/39 | 9/27 | 0/4 |

We next evaluated the effect of two glycosaminoglycans, heparan sulfate (HS) and heparin, on LECT2 fibrillogenesis by shaking. HS proteoglycans are expressed on the surfaces of endothelial and other cells and HS is known to bind hundreds of proteins^39^. HS has been detected in amyloid deposits of serum protein A, immunoglobin light chain, islet amyloid polypeptide, transthyretin, β2-microglobulin, prion protein, Alzheimer Aβ, and tau^40, 41^. Heparin, a potent anticoagulant, is mainly found in mast cells where it is sequestered from circulating proteins, but it has been widely used as a surrogate for HS to drive fibrillogenesis of many proteins^42^. HS enhanced amyloid formation for both I40V and WT apoLECT2 (Supporting Figure S3B), increasing the ThT fluorescence of 87 % of I40V and 18 % of WT samples over the course of the 200 h shaking experiments (Table 2). HS also globally decreased the lag times while maintaining the shorter values of I40V (compared to WT) observed in the absence of HS (Supporting Figure S12B).

Heparin promoted aggregation still further, causing 100 % of I40V and 33 % of WT samples to aggregate within 100 h, and shortening the lag times of both I40V and WT (<40 h in most cases) to the point where they became similar (Table 1 and Supporting Figure S12C). Glycosaminoglycans thus drive fibrillogenesis of apoLECT2, and the data further support the trend that the effect of the I40V mutation becomes masked when lag times are short. ThT fluorescence remained at baseline for all I40V and WT holoLECT2 controls whether they contained HS, heparin, or no additive (Supporting Figure S12D), indicating that zinc loss is required for aggregation in the presence as well as absence of glycosaminoglycans.

The data clearly reveal that flow shear induces misfolding and aggregation much more efficiently than shaking or stirring, at the same apoLECT2 concentration and temperature. Deposits of I40V apoLECT2 were visible in the channels within 24 h at 35 °C (Figure 2), whereas it generally took ∼100 h for aggregation to be detected in shaking experiments. This difference was even more pronounced with WT apoLECT2, which aggregated by flow shear (Figure 2 and Figure 3) but not by shaking (Table 2 and Supporting Figure S12A).

### EM structural analysis

We turned to TEM to determine whether the apoLECT2 particles formed in the microfluidic device were fibrillar. TEM samples were prepared by separating PDMF blocks from the coverslips, collecting the insoluble material by rinsing with water, and spotting the liquid onto grids. Negative stain TEM images revealed that apoLECT2 aggregates consisted primarily of large, darkly stained clumps with a minority of fields containing additional short, rod-shaped species (Figure 5A). Most of the clumps were too dense to make out detailed structural features; however, inspecting the periphery of some revealed fibrils that appeared to be in the process of joining or breaking off. No obvious difference between I40V and WT was observed for either the fibrils or the dense bodies. We therefore conclude that microfluidic flow caused I40V and WT apoLECT2 to aggregate into large, dense clumps that were composed of amyloid fibrils.

**Figure 5.**
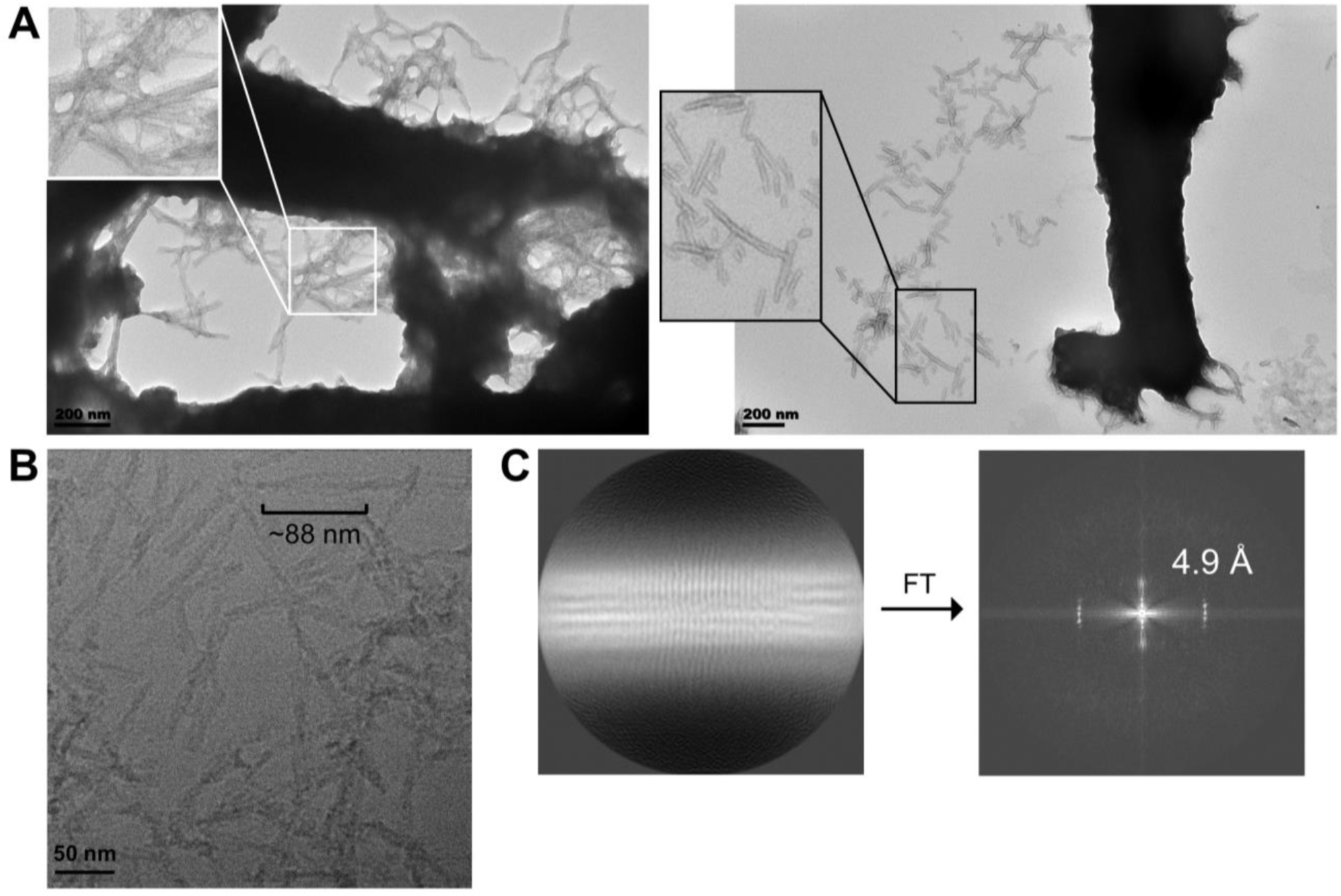
EM analysis of I40V apoLECT2 aggregates generated by flow shear using the microfluidic device (A) and by general shaking (B, C). **(A)** Aggregates recovered from the microfluidic channels were mainly large, dense particles composed of smooth, untwisted fibrils as determined by negative stain TEM. **(B)** Cryo-EM analysis of shaken samples revealed mostly short fibrils, with some of these exhibiting a helical crossover of ∼88 nm. **(C)** The power spectrum calculated from the 2D average of the fibrils shown in panel B indicated peaks at ∼(4.9 Å)^-1^.

Although flow shear and conventional stirring/shaking both induced apoLECT2 to form amyloid, two lines of evidence suggested the two methodologies generated distinct species. First, nearly all microfluidic aggregates consisted of large clumps of densely packed fibrils (Figure 5A) as opposed to the loosely associated and individual fibrils that resulted from stirring and shaking^7^ (Figure 5B). No particles of any size were visible by TEM in the supernatants of the microfluidic reservoirs after ultracentrifugation (200,000 x*g*) or even low-speed centrifugation (14,000 x*g*). By contrast, stirring and shaking produced significant quantities of individual, soluble fibrils as evidenced by the negative stain^7^ and cryo-EM^43^ (Figure 5B) images obtained from particles in the 10,000 – 14,000 x*g* centrifugation supernatants. Moreover, no pellets were recovered upon ultracentrifuging the microfluidic reservoirs, indicating that all aggregates were retained within the device in the form of the dense clumps shown in Figure 5A.

The second distinction was that most of the microfluidic fibrils appeared smooth (Figure 5A) whereas most of the fibrils generated by stirring and shaking exhibited pronounced helical twist. The tendency of the microfluidic fibrils to self-associate precluded cryo-EM analysis. However, cryo-EM images of shaken apoLECT2-tag fibrils revealed mostly individual fibrils, with some showing helical crossover of ∼88 nm (Figure 5B). Power spectra of individual fibers showed peaks at ∼(4.9 Å)^-1^, characteristic of the stacking of β-strands in amyloid fibrils^44^, which became more obvious in the 2D average generated from a small dataset of helical segments processed in RELION-4^45^ (Figure 5C). A recent cryo-EM study obtained a similar result with apoLECT2 amyloid induced by rapid shaking, reporting an 85 nm helical crossover for 90 % of fibrils and a 40 nm crossover for the remaining 10 %^43^. These data suggest that fluid shear generates smooth, ‘sticky’ fibrils that associated into large clumps more readily than fibrils created by stirring and shaking. The network of branched channels essentially filtered these clumps and became clogged, reminiscent of kidney failure in late stage ALECT2.

### Flow shear generated misfolded protein that was capable of seeding

One of the hallmarks of pathogenic amyloid is its ability to nucleate formation of new fibrils from monomers. Potential nuclei were obtained by collecting the reservoirs at the conclusion of the I40V experiments shown in Figure 2 and ultracentrifuging them to remove insoluble particles. As noted above, no protein was detected in the pellets, and Trp absorbance scans of the supernatants showed only a small reduction in protein concentration (compared to controls that had not been circulated through the microfluidic device; Supporting Figure S13), commensurate with the amyloid that had accumulated in the chip. The only indication of potential nuclei was the small amount of light scatter in the absorbance scans which suggested soluble oligomers large enough to scatter ultraviolet light.

Seeding was performed by mixing 1 ml of the above supernatant with 1 ml of fresh I40V apoLECT2. The unseeded controls consisted of 2 ml of fresh I40V apoLECT2 run side-by-side on the same chip. Analyzing the images as in Figure 2 found that the area and total intensity of aggregates were greater in the seeded sample at all times examined (Figure 4B and Figure 6; see also Supporting Figure S14). The average intensity per pixel also tended to be higher, but the difference was not always significant. Thus, while I40V aggregates were consistently denser than WT aggregates (Figure 2), nucleating I40V monomers with I40V nuclei did not increase density of the resulting particles to the same extent. This result may be anticipated since seeding is expected to increase the rate of amyloid formation but not necessarily change the properties of the final structures.

**Figure 6.**
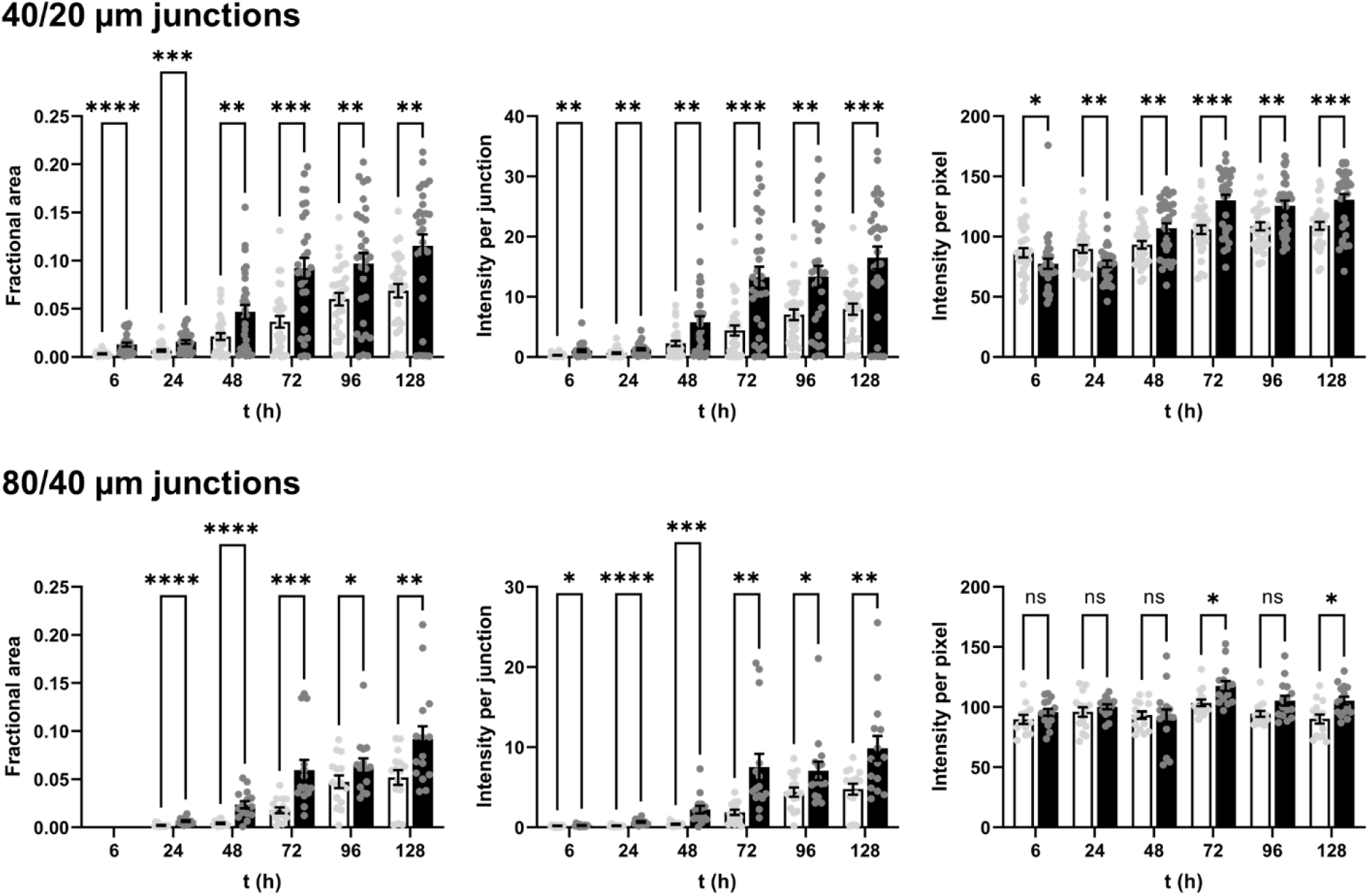
Misfolded I40V apoLECT2 nucleated aggregation of I40V apoLECT2 monomers. Light and dark bars indicate unseeded and seeded samples, respectively. Number of data points and analysis method are identical to those in Figure 2. ****, p<10^-4^; ***, p<10^-3^; **, p<10^-2^; *, p<10^-1^; ns, not significant. See Supporting Figure S14 for additional BR data.

## Discussion

This study established for the first time that the I40V mutation enhances fibrillization of LECT2. Loss of the single bound zinc ion was essential to this process. We have therefore identified two physiologically reasonable conditions that may combine to cause ALECT2. Using the data in Table 1 and the concentration of available Zn^2+^ in the blood (≈10 nM)^46^, 2 % of LECT2 is predicted to exist in the apo form at physiological pH of 7.4. Zinc loss is exacerbated by acidic conditions (affinity drops by 60-fold at pH 6.5 due to protonation of the zinc-ligating His residues)^7^ and conceivably by low dietary zinc intake, a condition associated with elderly Hispanics^47^ who comprise the U.S. demographic most afflicted with ALECT2. Heparin and the ubiquitous polysaccharide HS also facilitated aggregation.

The second notable finding of this work is that kidney-like flow shear induces apoLECT2 aggregation much more efficiently than traditional methods of stirring or shaking. Extensive amyloid deposits were observed in the microfluidic channels within 24 h of pumping at a flow rate of 10 µl/min. By contrast, aggregation required many days of vigorous shaking with silica beads, and even then, two-thirds of I40V samples and all WT samples failed to form amyloid. Reproducing the branched networks in the kidney appeared to be important, as aggregation began at the smallest, terminal junctions and then proceeded upstream.

The efficiency of flow shear in inducing misfolding and subsequent fibrillogenesis is remarkable considering that, with typical reservoir volumes of 2 – 6 ml, the average protein passed through the device only 2.4 – 7.2 times per 24 h. These are far fewer cycles than would occur in the human kidney, which processes all 4.5 L of one’s blood almost twice per hour. Our data indicate that only a few transits through kidney-sized, branched channels was sufficient to cause apoLECT2 to misfold. Aggregation followed rapidly—much more so than in ALECT2 disease which usually manifests in older adults—presumably because the LECT2 concentration in our experiments (10 µM) was ∼10^4^-fold higher than that present in the blood^48, 49^. Regarding the latter point, it is noteworthy that the amyloid-inducing effect of the I40V mutation grew more pronounced as aggregation became more gradual. It seems reasonable to speculate that the lower concentrations of LECT2 in the blood would both slow amyloid formation and make it more conditional on the presence of the I40V mutation, compared to the data we presented here.

Our results add to the growing body of evidence that different forms of mechanical disruption produce different amyloid structures and highlight the importance of inducing protein misfolding in a manner consistent with the disease process^50^. To this end, Knowles and co-workers emphasized the importance of fluid flow in confined, microscale channels to reproduce the laminar conditions usually present in blood vessels^31^. The resulting shear stretches and distorts proteins, exposing potentially amyloidogenic sequences^29^. This type of disruption is distinct from that achieved by methods such as atomic force microscopy (which pulls proteins at specific points of attachment) and air-water interface adsorption in which proteins restructure to orient their buried hydrophobic residues toward the air layer. In the latter case, most of the α-helices in lysozyme transformed into β-strands^51, 52^, a mechanism of fibrillogenesis that is likely potent but artificial.

## Conclusions and implications

The results suggest that devices such as ours may be of general use to study systemic amyloidoses involving misfolding of circulating proteins (*e.g*., immunoglobin light chain, serum amyloid A, transthyretin), of which ∼30 examples have been reported to date. All circulating proteins experience the same kidney flow shear-induced stress that we attempted to reproduce here. The effects of hypertension (another condition that affects the U.S. population with the highest incidence of ALECT2^2, 53^) on protein misfolding and aggregation can be modeled by our device by modulating flow rate and pressure. Given the current interest in characterizing patient-derived amyloid, and the finding that their structures and properties can differ from aggregates generated by stirring and shaking^50, 54–56^, it is important to develop methods for recapitulating the physiological process of amyloidosis *in vitro* so that its molecular mechanisms can be delineated and inhibitors can be developed. The device created here can be fabricated in large quantities and inexpensively, opening the door to the above studies.

## Methods

### Gene construction, protein expression, and protein purification

The LECT2 gene (without the region encoding the N-terminal signal sequence) was placed at the 3’ end of the *Thermoanaerobactor tencongensis* ribose binding protein (RBP) gene, using a sequence encoding for 25 amino acid linker that included an HRV3C protease recognition site. A sequence encoding a His6-tag was placed at the 5’ end of the RBP gene. Proteins were expressed in *E. coli* and purified as previously described^7^, whereupon RBP was cleaved away and LECT2 was purified to >98 % homogeneity using a Superdex S75 column. LECT2 concentration was determined by absorbance (ɛ_280_ = 15,010 M^-1^ cm^-1^; MW = 14,798 Da).

### Protein stability and zinc binding

Experimental conditions were 2.5 µM protein, 20 mM Tris (pH 7.5), 0.15 M NaCl, 37 °C. GdnHCl denaturation experiments were performed as previously described^7^ except 1 mM EDTA was added to the apoLECT2 samples and 7.5 µM ZnCl_2_ and 22.5 µM iminodiacetic acid was added to the holoLECT2 samples to buffer [Zn^2+^]_free_ at a constant concentration of 6-8 µM. Data were fit to the two-state linear extrapolation equation^57^ to obtain thermodynamic parameters. Equilibrium zinc binding and zinc dissociation kinetic experiments were performed as described previously^7^.

### Microfluidic experiments

Microfluidic devices were fabricated at the Cornell University NanoScale Facility using soft lithography methodology. The channel design was patterned onto a 5 x 5-inch chrome photomask which was then transferred to a 100 mm diameter n-type silicon wafer with photoresist by contact printing. This master pattern was plasma etched into secondary wafers to the desired depth and then treated with fluoroctatrichlorosilane to aid the peel-off process. When producing the microfluidic devices with the master wafer, a 1:10 mixture of Sylgard™ 184 crosslinker to elastomer was mixed and poured onto the silicon wafer. The wafer with the desired thickness of PDMS mixture on top was placed into a desiccator to degas for 30 min, then into a 110 °C oven to cure for 35 min, and finally cooled to room temperature on a benchtop. The PDMS block was cut out and inlet/outlet holes were punched. It was then bonded to a glass slide using a plasma cleaner (25 W RF power for 90 s) and placed in a 90°C oven for 1 h to stabilize the bonds. Microbore tubes were inserted into the pre-punched inlet/outlet holes and sealed off with a small amount of PDMS and the device was placed in a 60°C oven to cure without damaging the microbore tubing. Finally, the finished devices were cleaned by flowing in 70 % ethanol for 2 h followed by a 2 h flush with the buffer used in the experiments (below).

For aggregation experiments, reservoirs containing freshly thawed 10 µM I40V or WT apoLECT2 (in 20 mM Tris, pH 7.5, 0.15 M NaCl) were placed in a 35 °C temperature bath and circulated simultaneously through parallel channels on the same chip, using the same peristaltic pump to ensure consistency. Apo and holo LECT2 samples contained EDTA and ZnCl_2_/iminodiacetic acid, respectively, as described above. Experiments with glycosaminoglycans contained 5 μg/ml heparin or 5 μg/ml HS. Protein aggregation was monitored by recording images of the 160/80 μm, 80/40 μm, and 40/20 μm channel junctions using a Zeiss Axio Observer Z1 inverted microscope mounted with a 10x objective lens.

The data were first processed using Fiji^58^ by inverting the raw brightfield images and subtracting background with a 50-pixel sliding ball radius. The resulting images were then processed in MATLAB (MathWorks, Inc.). Images were aligned (using the *imregister* fuction) to a reference binary image that contained empty channels. We wrote a custom code to manually correct misalignments using *imtranslate* and *imrotate* functions. The channels were then marked with a custom mask that was made from the reference image. The aggregates were identified by adjusting the threshold intensity according to Otsu’s method using the *imbinarize* function, then picking the pixels that also contained the mask. The fractional area of aggregates was calculated by dividing the number of pixels that contained aggregates by the total number of pixels in the channels. The integrated intensity per junction was calculated by measuring the intensity of the pixels in the channels that contained aggregates, subtracting the average intensity of the pixels in the channels that did not contain aggregates, and dividing that value by the number of pixels per channel.

Computational fluid dynamic calculations were performed using COMSOL Multiphysics® to simulate flow in the microfluidic device and quantify flow velocity, shear rate, strain rate, and Reynolds number.

### Shaking experiments

150 µL of freshly thawed apoLECT2 (same concentration and buffer conditions as in the microfluidic experiments) was added to wells of fully blackened, polystyrene, round bottom 96-well plates (Corning CoStar). Six 1-mm diameter Zirconia/silica beads (BioSpec) and 15 µM ThT were added to each well and the plates were sealed (TempPlate RT Select Optical Film; USA Scientific) to prevent evaporation. Plates were shaken at 650 RPM and 34 °C using a Mixer HC (USA Scientific). At the indicated times, ThT fluorescence was recorded using a SpectraMax i3x plate reader (Molecular Devices) with excitation at 410 nm and emission at 490 nm.

### Electron microscopy

Aggregated proteins from the microfluidic device were obtained by excising the PDMS block from the glass slides using a sterile scalpel and flushing the surfaces repeatedly with 100 µl of ultrafiltered deionized water. The collected liquid was refrigerated along with the remaining liquid in the reservoirs until further analysis. Procedures for collecting aggregated proteins from shaking experiments as well as for EM sample preparation were described previously^7^. Negative stain EM data were collected on a JEOL JEM1400 electron microscope with a Gatan Orius 832 CCD. Cryo-EM data were collected on a JEOL JEM-2100F microscope with K2 Summit direct detector. Data were processed using EMAN2^59^ and RELION-4^45^ software packages.

### Data sharing statement

All data generated in this study, including raw images of aggregates, calculated areas and intensities of aggregates, and code generated for processing images, have been deposited to the Figshare Data Repository (DOI 10.58120/upstate.23660583).

## Author contributions

S.N.L., J-H.H., and D.R. conceived the work. Y.X. and D.R. designed the microfluidic device. Y.X. performed the microfluidic experiments, J-H.H. and S.W. performed TEM studies, and J-H.H. carried out all other experiments. J-H.H. and H.S. analyzed the microfluidic data using the methods developed by H.S. S.N.L. and J-H.H. wrote the manuscript with input from all authors.

## Acknowledgements

This work was funded by NIH grants R35 GM141908 to S.W., R01 EB030621 to D.R., F30 GM146428 to H.S., and R56 DK124335 to S.N.L. The Cornell NanoScale Facility is a member of the National Nanotechnology Coordinated Infrastructure (NNCI) and is supported by National Science Foundation grant NNCI-2025233.

## Supporting Information for

**Supporting Figure S1, related to Table 1.**
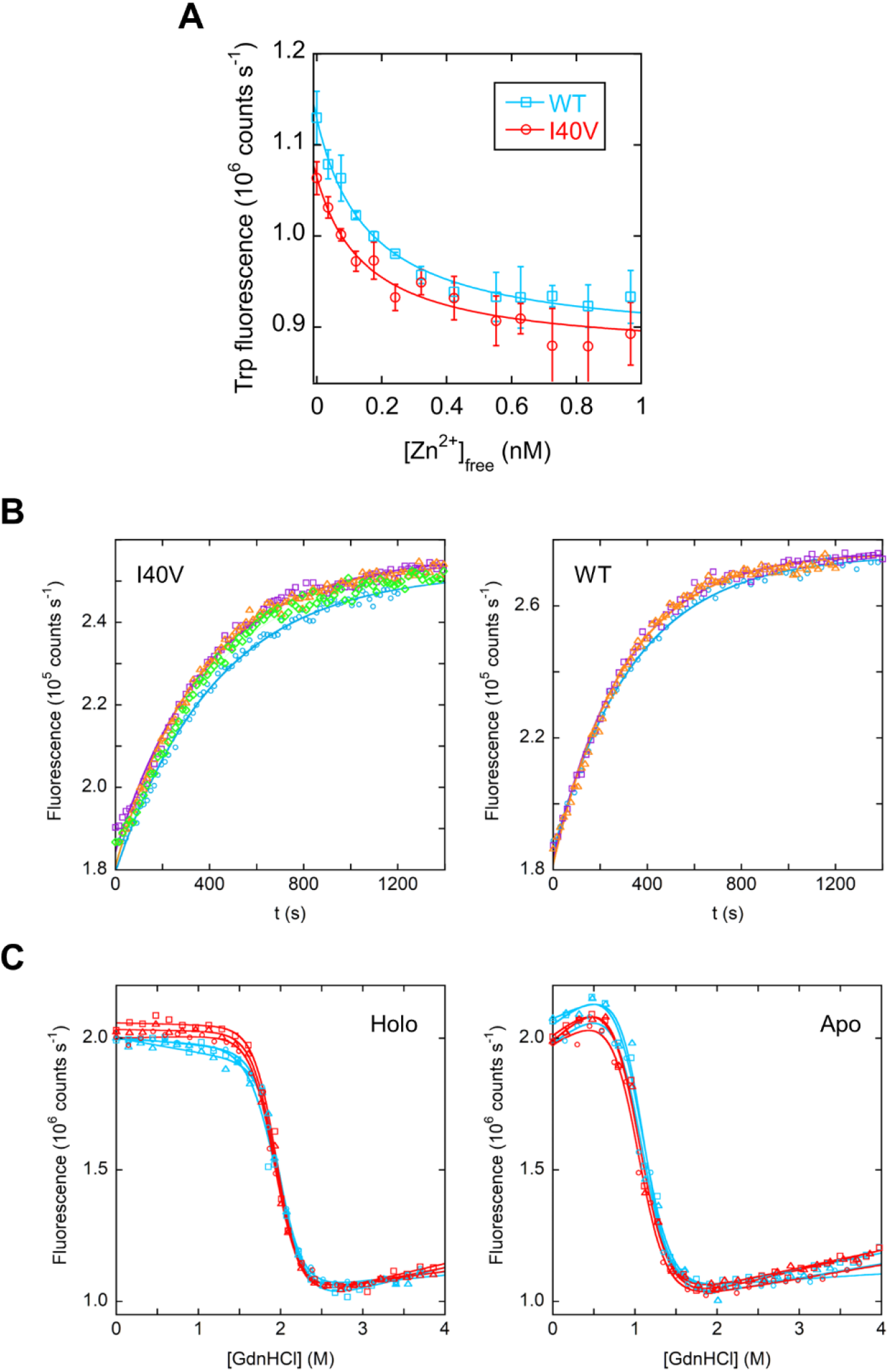
Zinc binding and stability of LECT2. **(A)** I40V and WT LECT2 bound zinc with comparable affinities. Lines are best fits to the quadratic binding equation. Error bars are s.d. (n = 3). **(B)** I40V and WT LECT2 exhibited similar off-rates. Lines are best fits to a single exponential function. **(C)** The I40V mutation selectively destabilized the apo form of LECT2 compared to the holo form. Red and blue data indicate I40V and WT proteins, respectively. Lines are best fits to the linear extrapolation equation for two-state denaturation.

**Supporting Figure S2, related to Figure 1.**
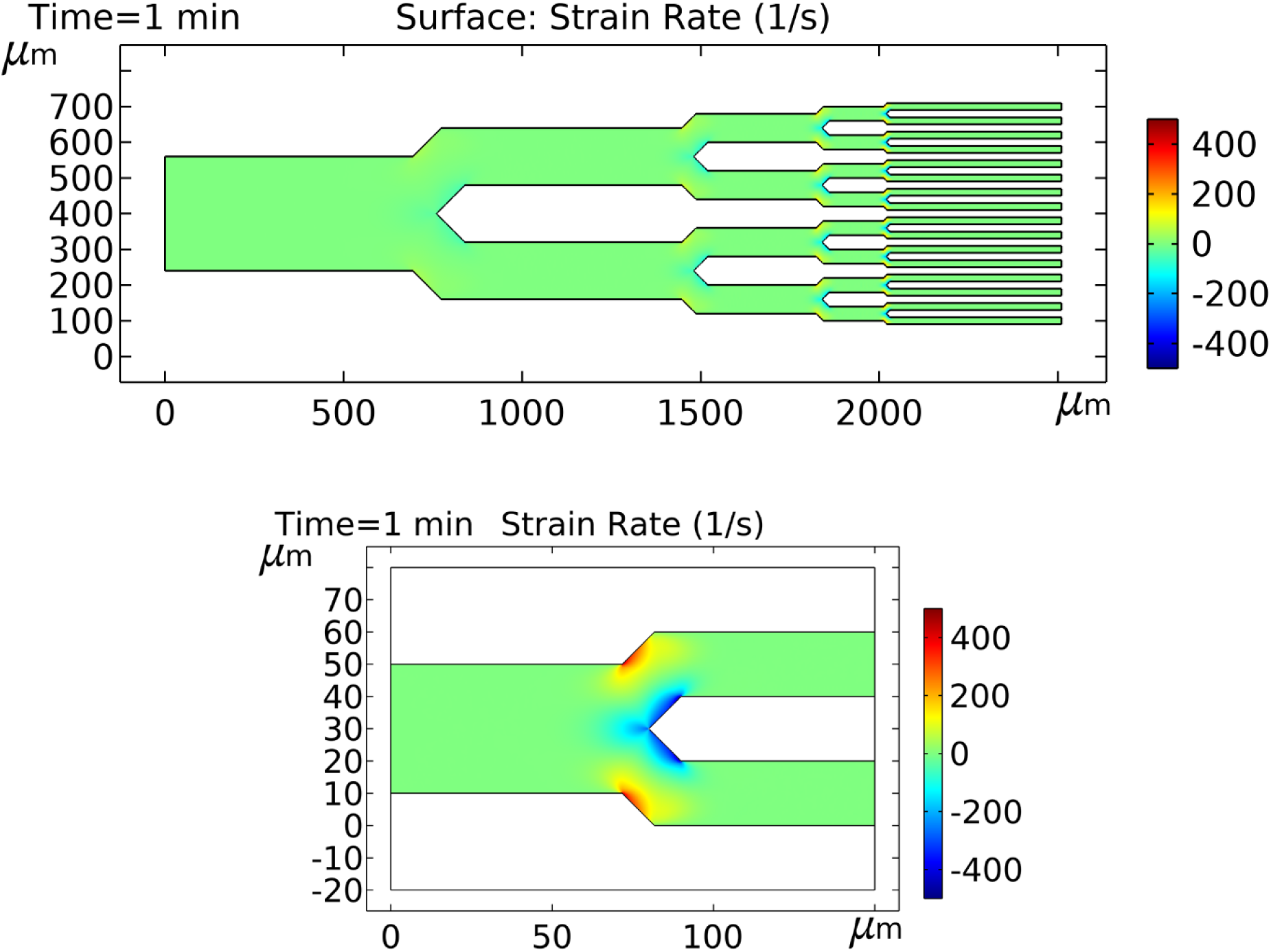
Compared to flow shear, flow strain was minimal in the microfluidic device. COMSOL was used to calculate strain rates. The 320 µm – 20 µm channel path is shown at top, with the 40/20 µm junction expanded at bottom.

**Supporting Figure S3, related to Figure 2.**
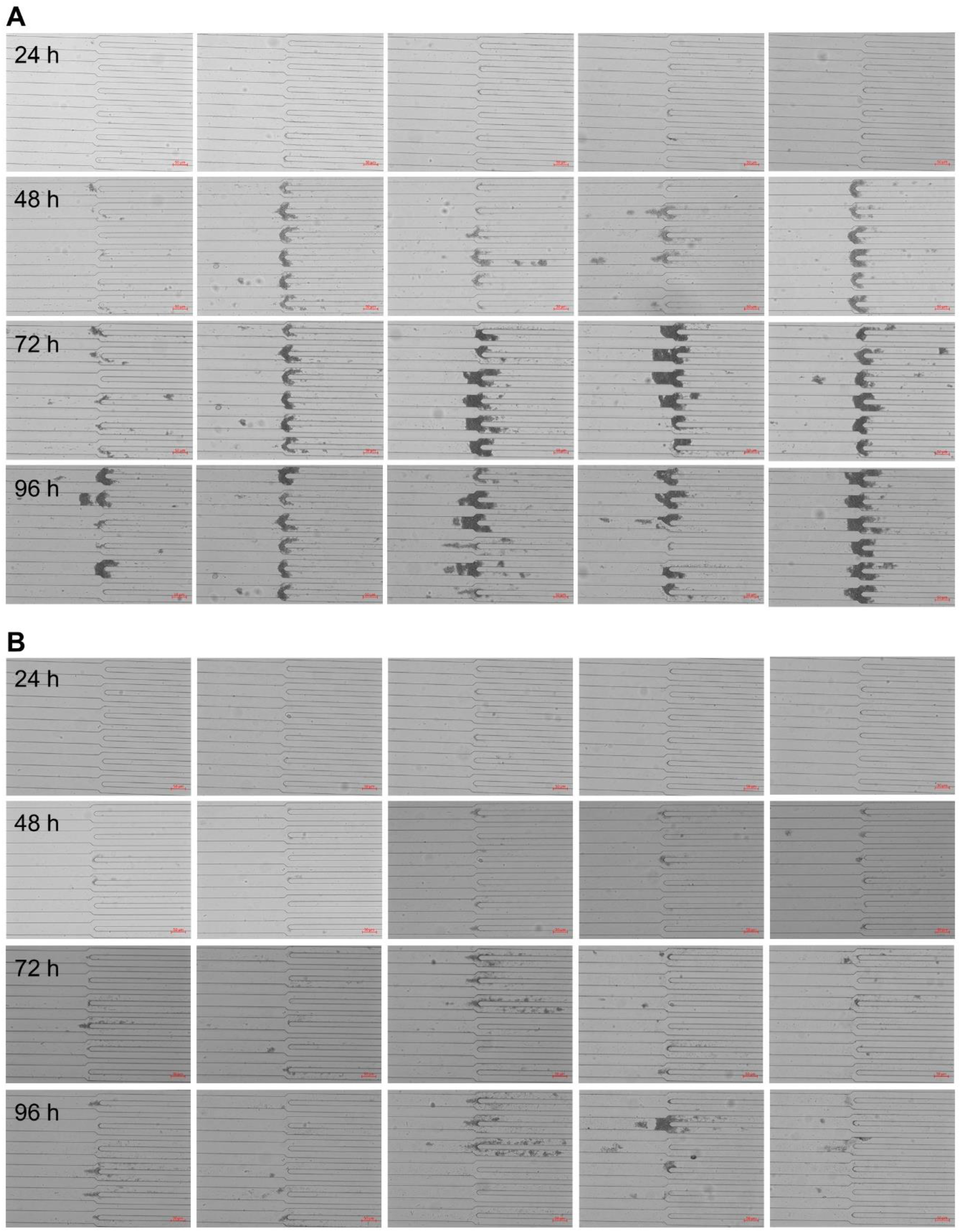
Images of 40/20 µm junctions from the microfluidic chip, obtained from a second BR. I40V and WT apoLECT2 are shown in **(A)** and **(B)**, respectively.

**Supporting Figure S4, related to Figure 2.**
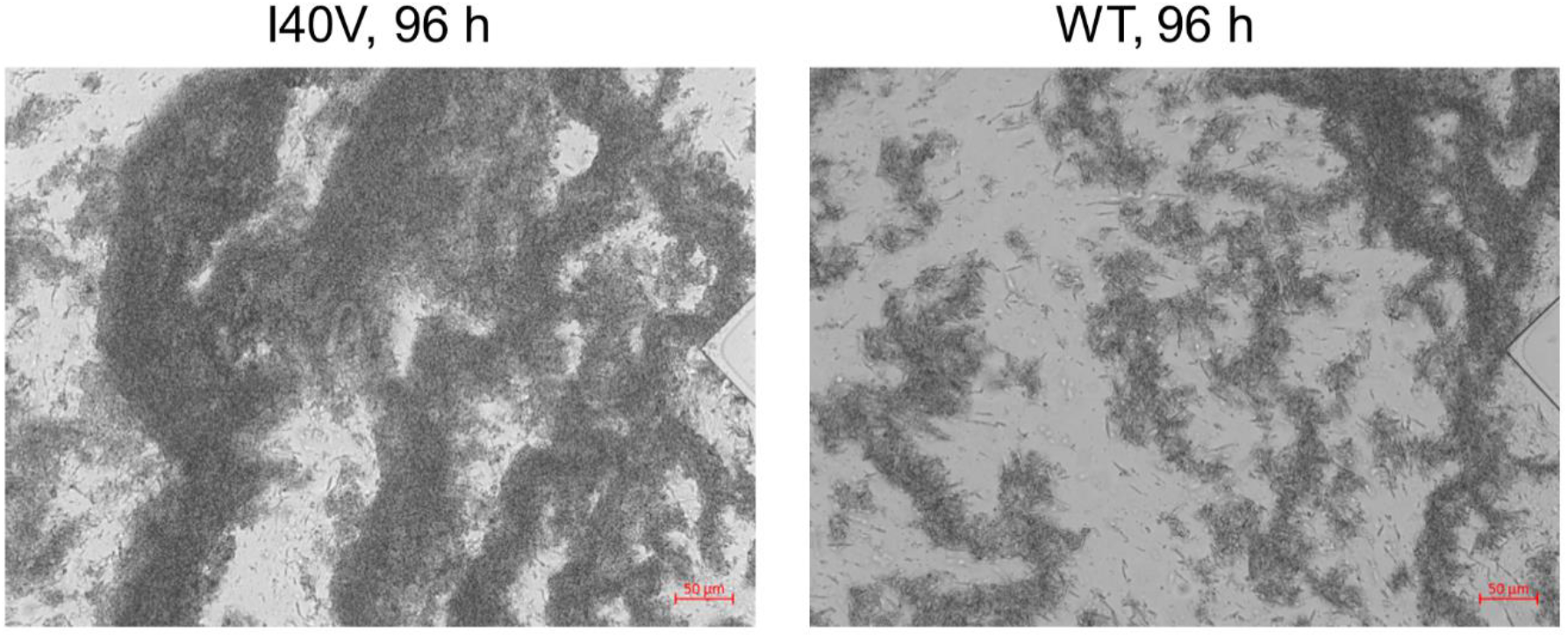
Images of 1280/640 µm junctions from the microfluidic chip. Scale bars are 50 µm.

**Supporting Figure S5, related to Figure 3.**
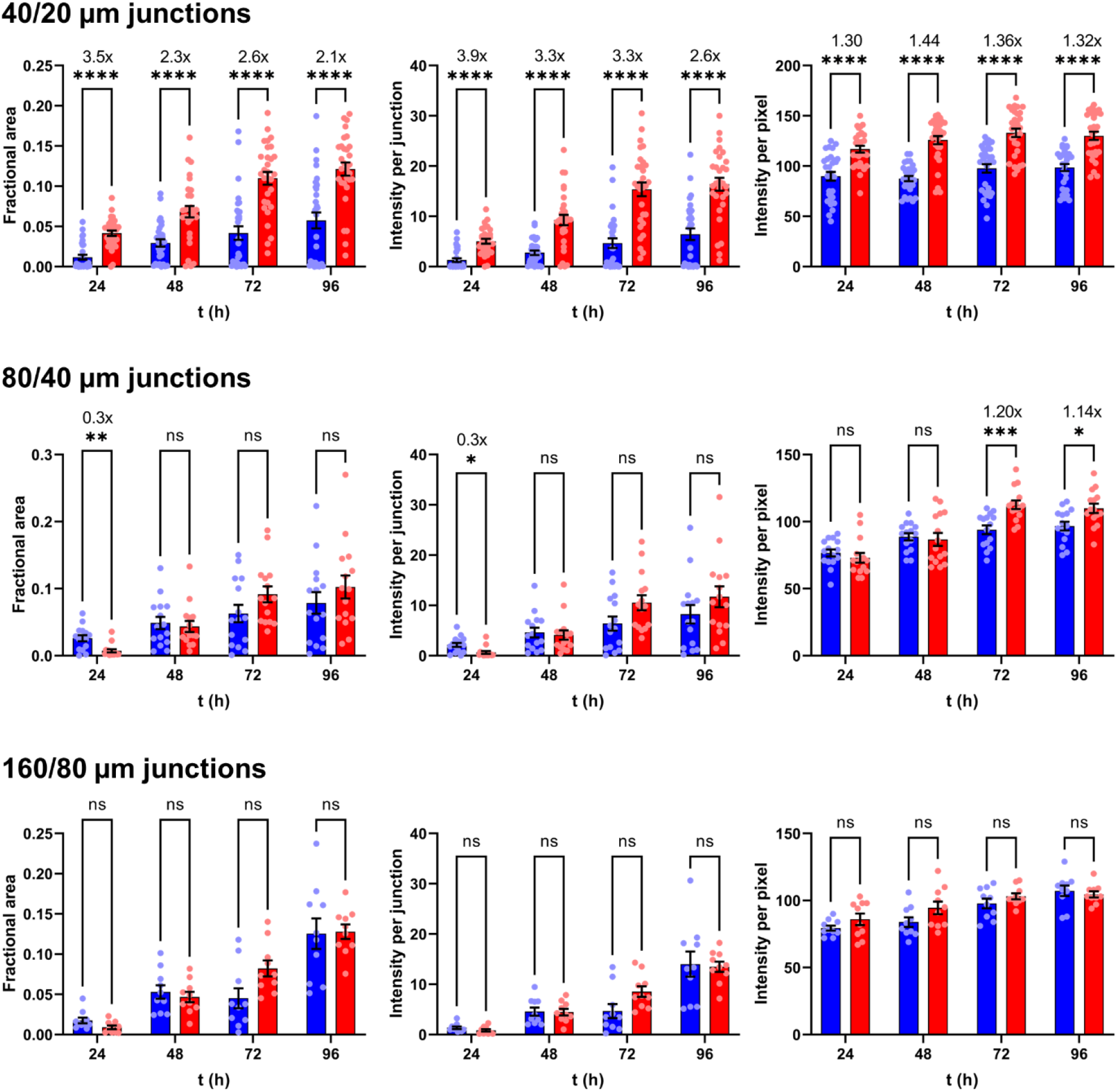
Quantification of microfluidic aggregates from an additional BR. Red and blue data indicate I40V and WT apoLECT2, respectively. See Figure 3 for description of statistical analysis.

**Supporting Figure S6, related to Figure 3.**
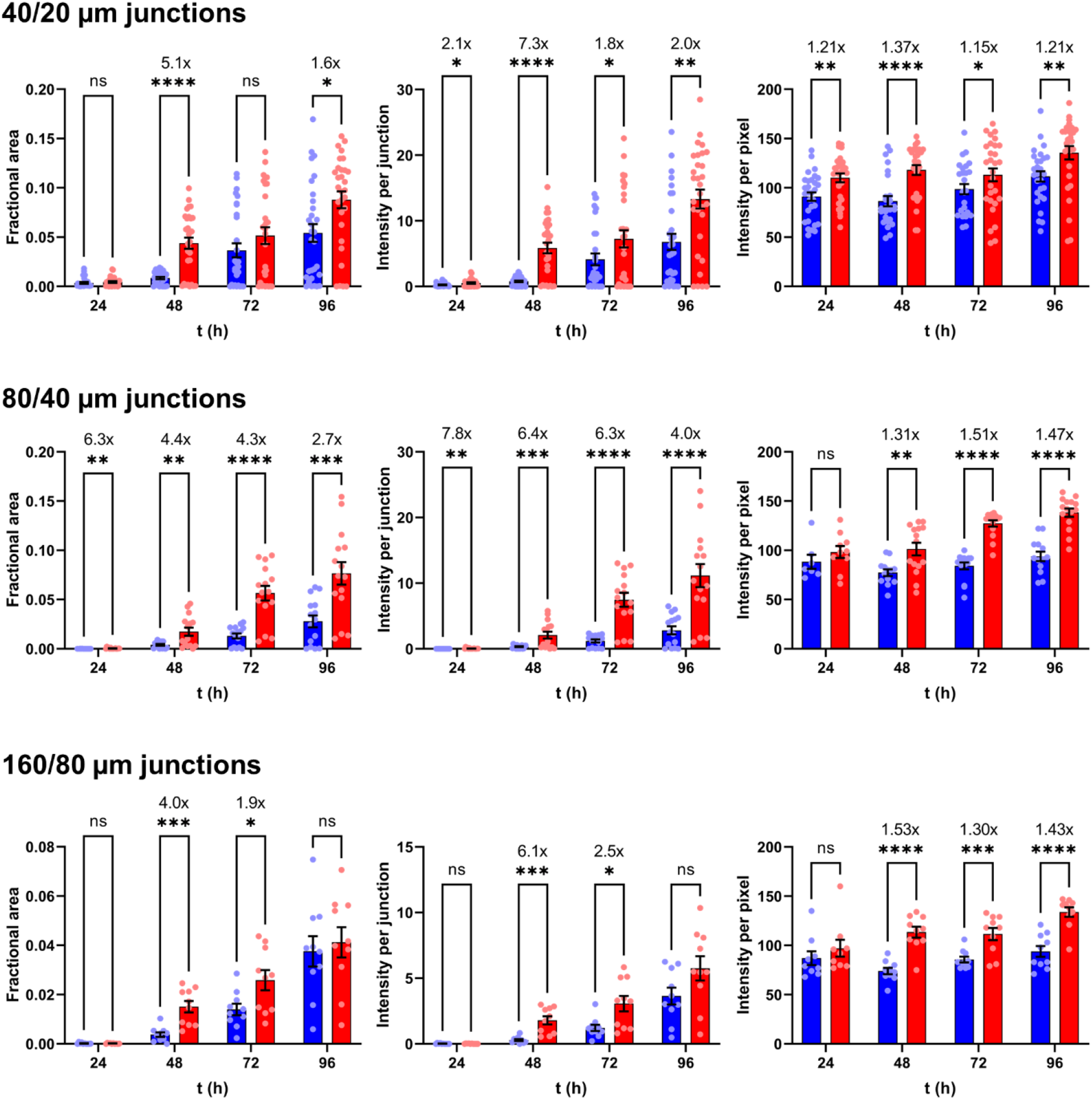
Quantification of microfluidic aggregates from an additional BR. Red and blue data indicate I40V and WT apoLECT2, respectively. See Figure 3 for description of statistical analysis.

**Supporting Figure S7, related to Figure 3.**
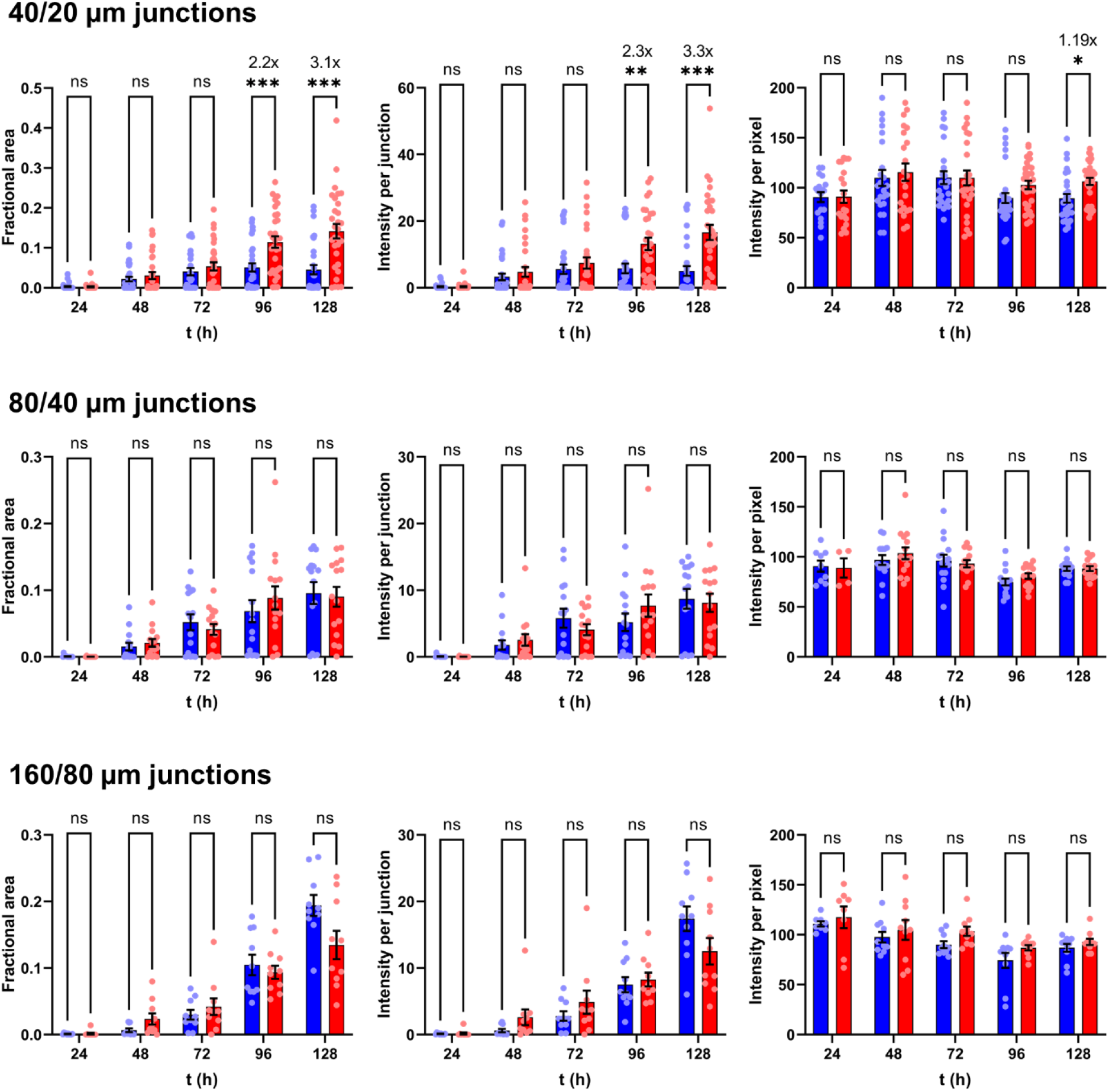
Quantification of microfluidic aggregates from an additional BR. Red and blue data indicate I40V and WT apoLECT2, respectively. See Figure 3 for description of statistical analysis.

**Supporting Figure S8, related to Figure 3.**
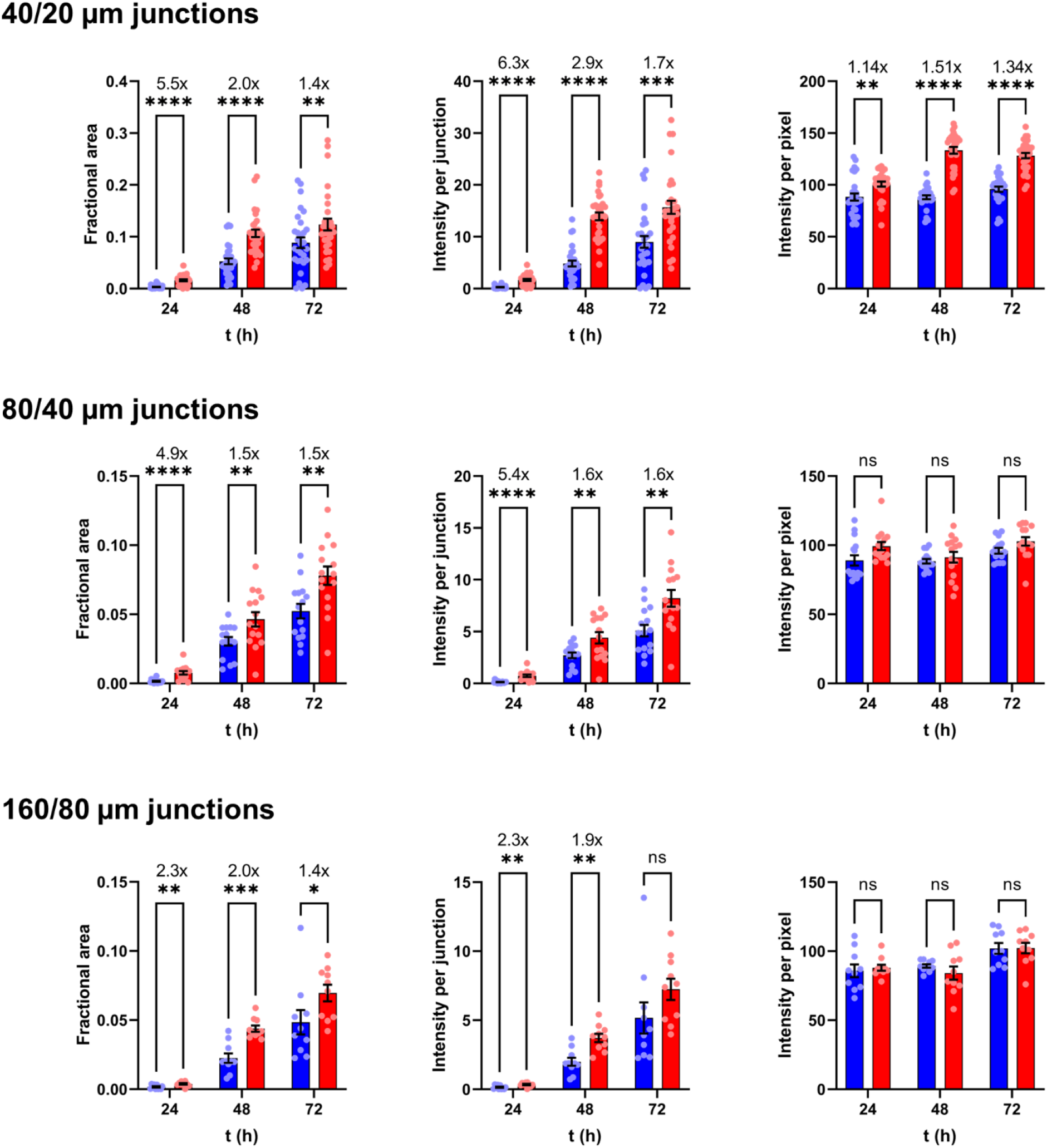
Quantification of microfluidic aggregates from an additional BR. Red and blue data indicate I40V and WT apoLECT2, respectively. See Figure 3 for description of statistical analysis.

**Supporting Figure S9, related to Figure 3.**
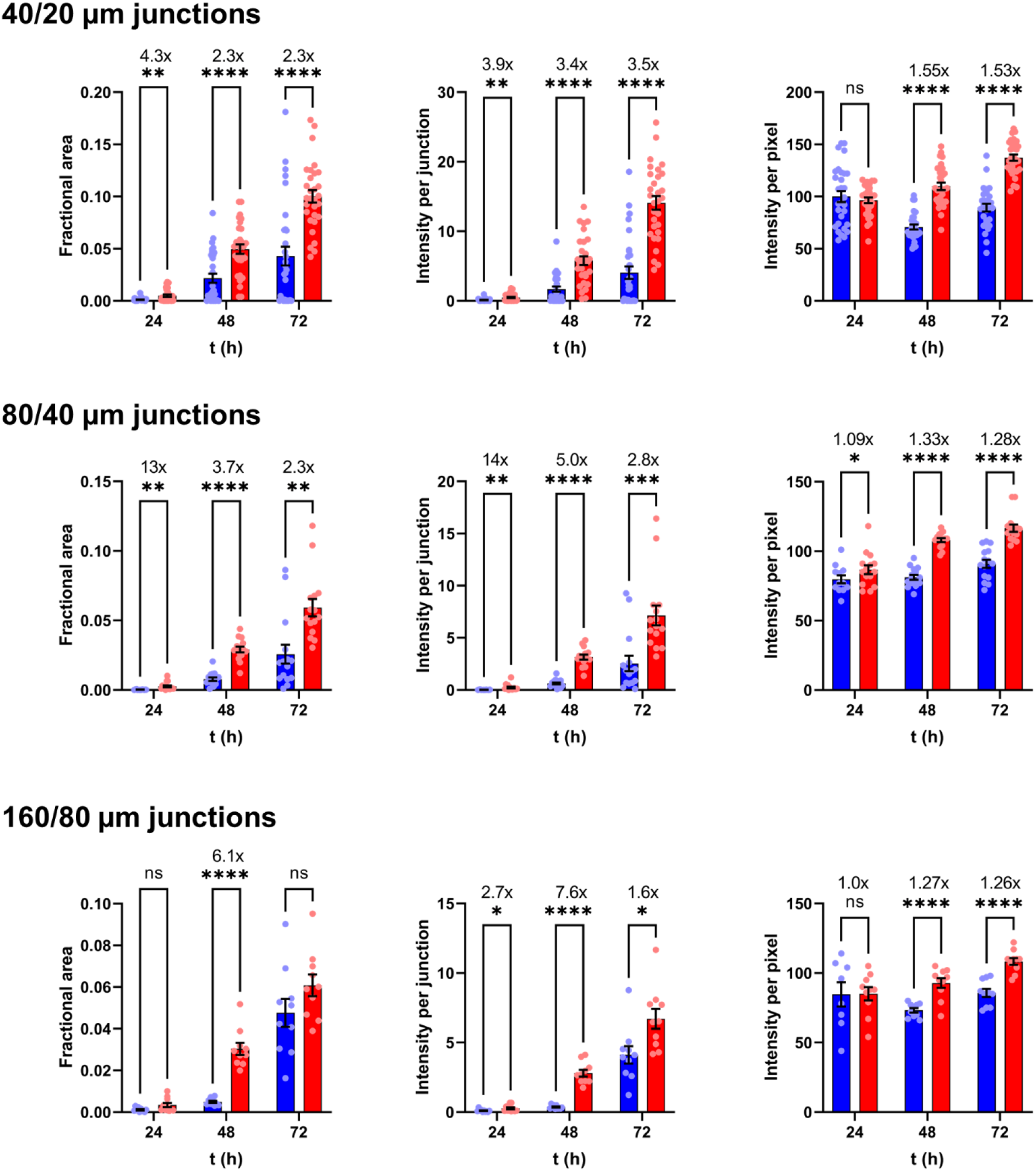
Quantification of microfluidic aggregates from an additional BR. Red and blue data indicate I40V and WT apoLECT2, respectively. See Figure 3 for description of statistical analysis.

**Supporting Figure S10, related to Figure 3.**
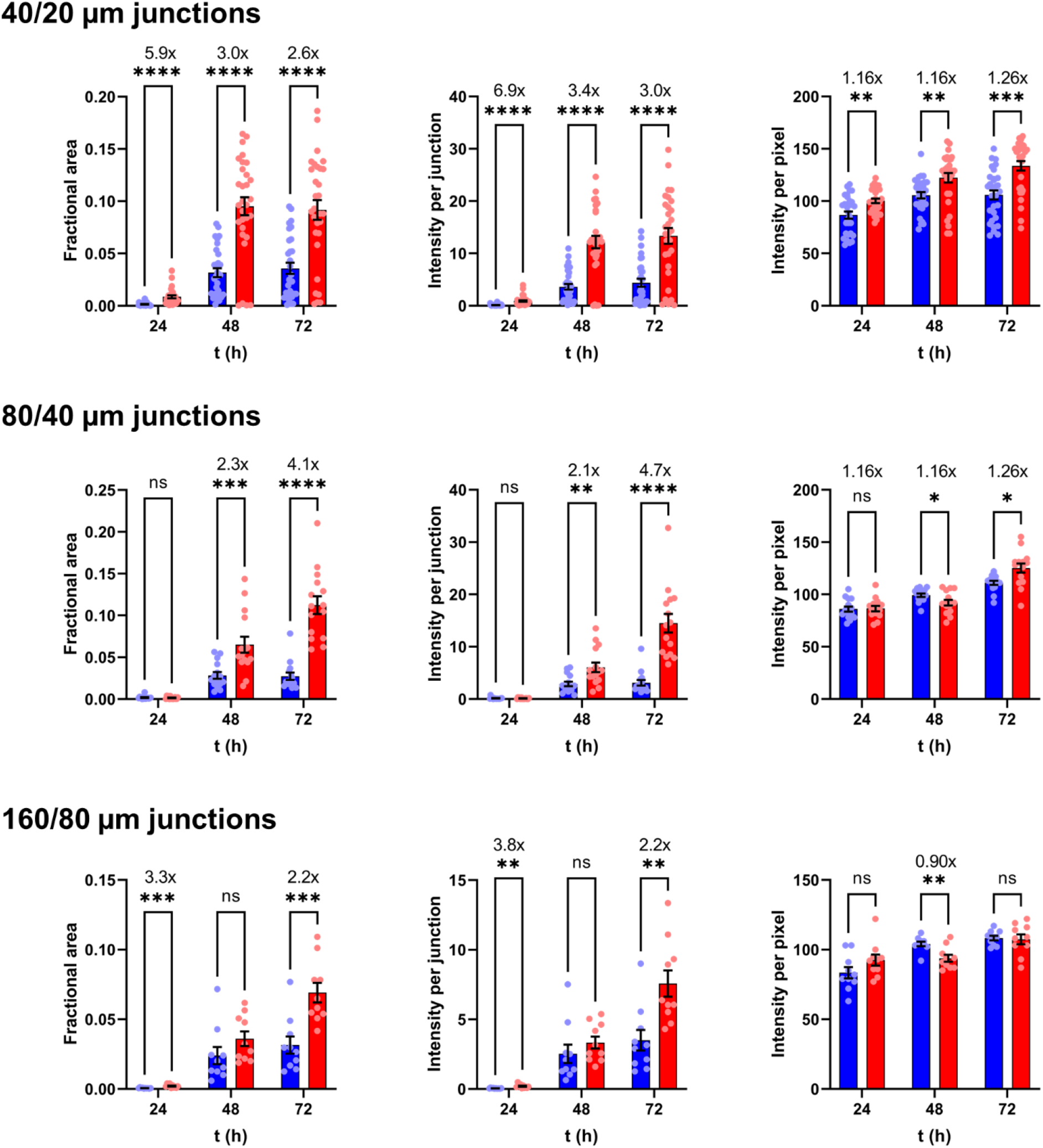
Quantification of microfluidic aggregates from an additional BR. Red and blue data indicate I40V and WT apoLECT2, respectively. See Figure 3 for description of statistical analysis.

**Supporting Figure S11, related to Figure 3.**
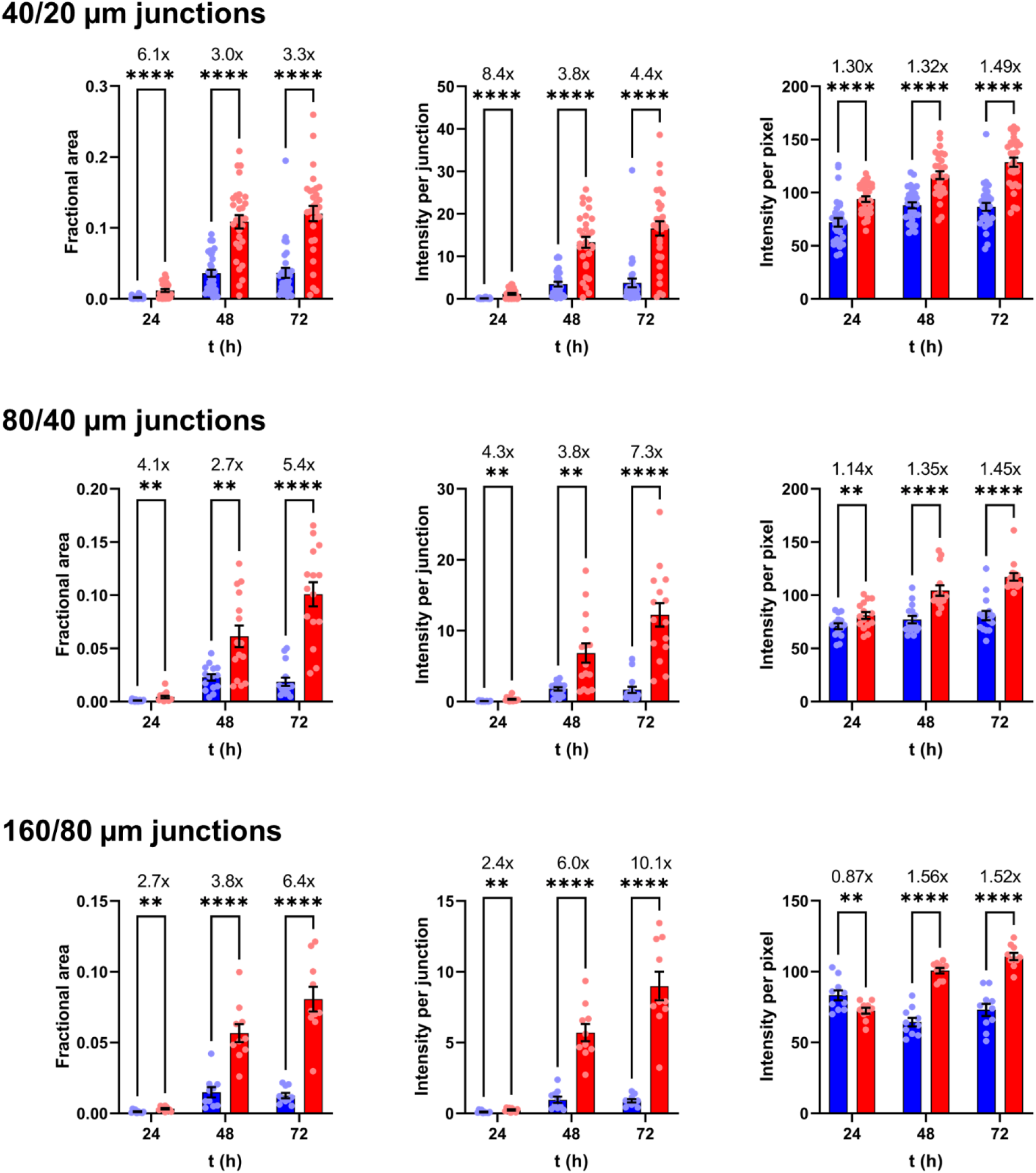
Quantification of microfluidic aggregates from an additional BR. Red and blue data indicate I40V and WT apoLECT2, respectively. See Figure 3 for description of statistical analysis.

**Supporting Figure S12, related to Table 2.**
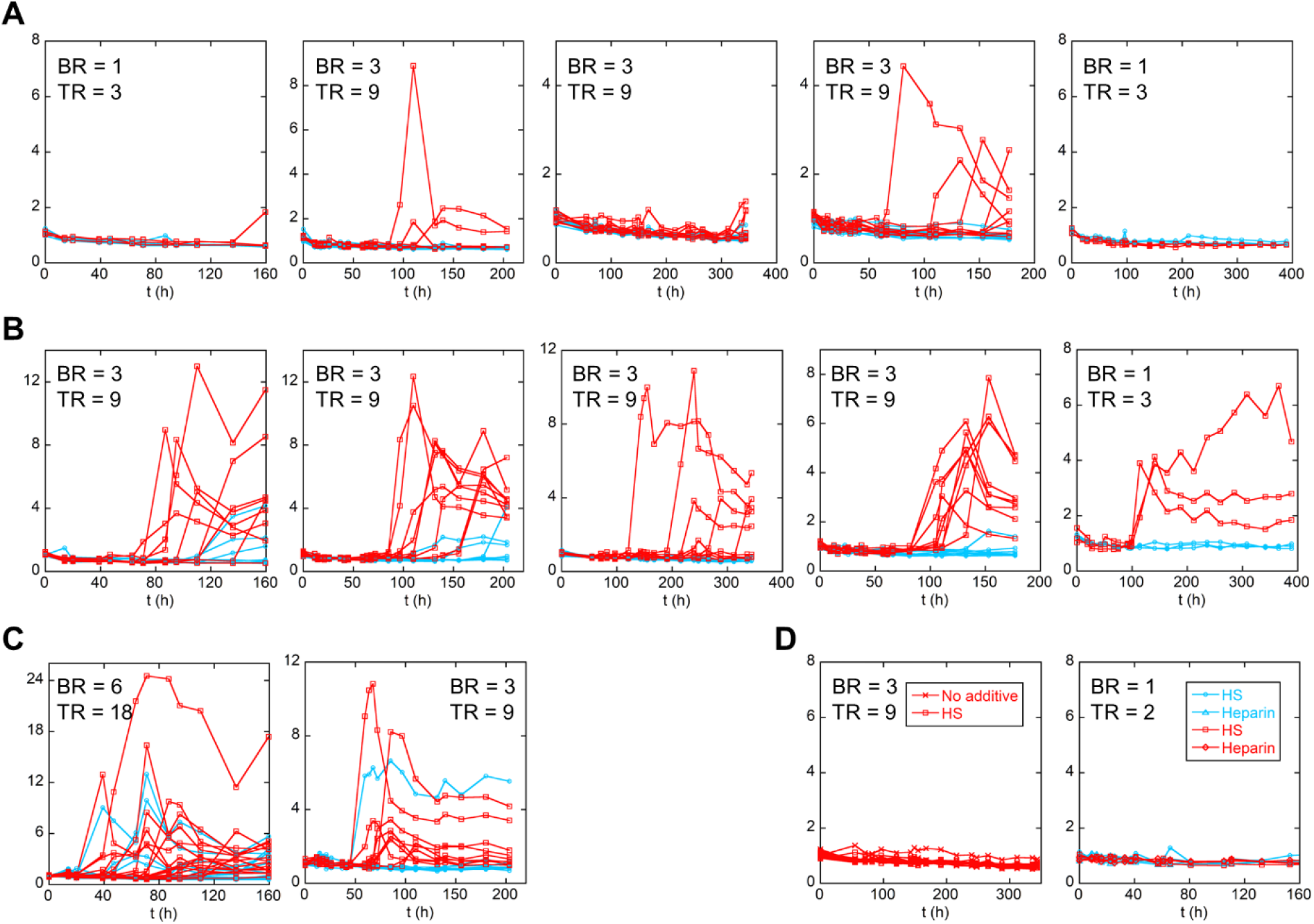
Shaking-induced fibrillogenesis of apoLECT2 monitored by ThT fluorescence. Data are shown for I40V apoLECT2 (red) and WT apoLECT2 (blue) with no additive **(A)**, HS added **(B)**, and heparin added **(C)**. Lines are meant to guide the eye only. **(D)** I40V holoLECT2 (red) and WT holoLECT2 (blue) controls did not form amyloid in the absence of additive, or with HS or heparin added.

**Supporting Figure S13, related to Figure 2 and Figure 6.**
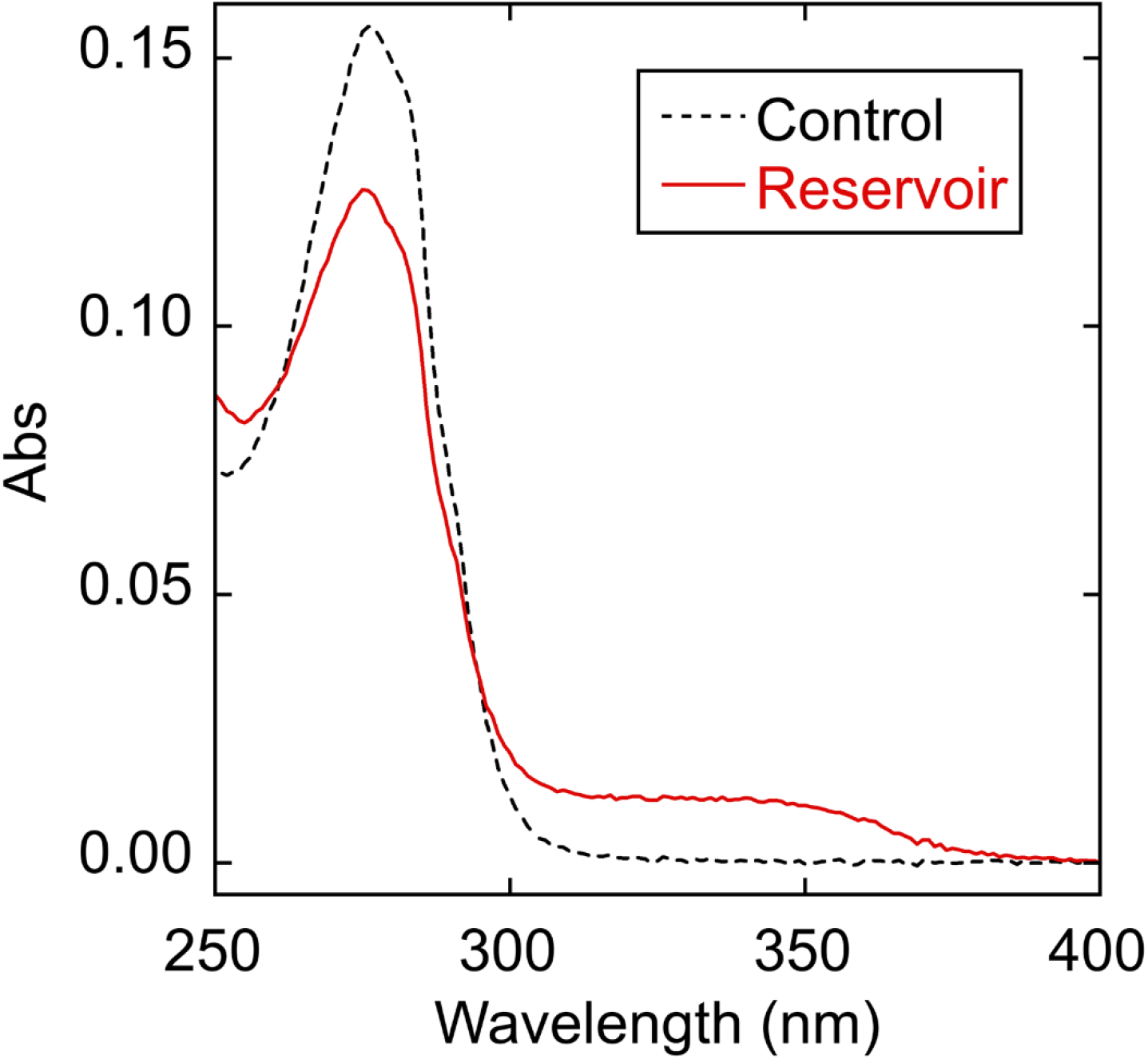
Absorbance scans of reservoirs from I40V apoLECT2 microfluidic experiments shown in Figure 2. The control spectrum is that of an identical sample that had not been passed through the device.

**Supporting Figure S14, related to Figure 6.**
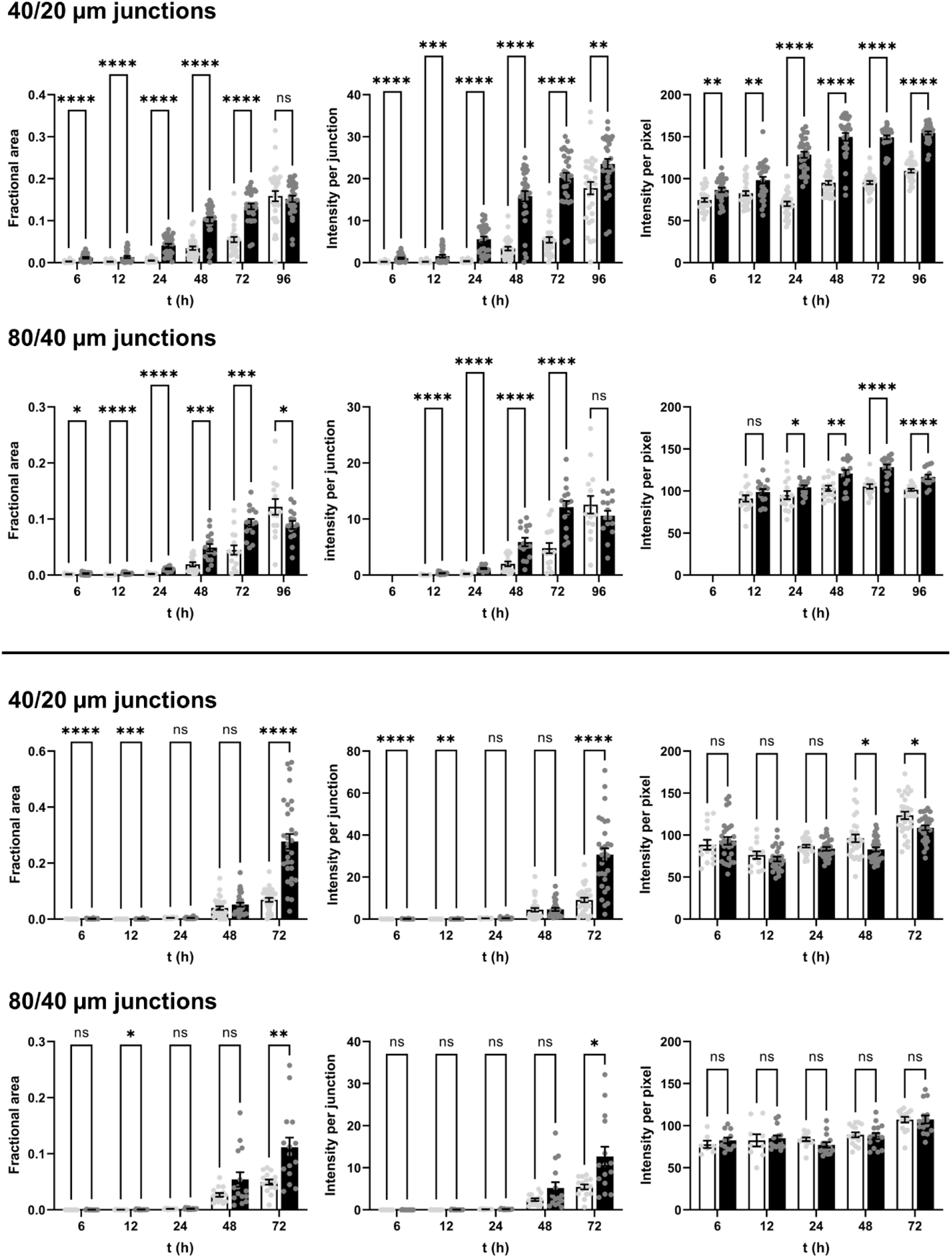
Quantification of microfluidic aggregates from two additional BRs. Black and white bars indicate seeded and unseeded (respectively) samples of I40V apoLECT2. See Figure 6 for description of statistical analysis.

